# A clade of RHH proteins ubiquitous in Sulfolobales and their viruses regulates cell cycle progression

**DOI:** 10.1101/2022.07.28.501860

**Authors:** X Li, C Lozano-Madueño, L Martínez-Alvarez, X Peng

## Abstract

Cell cycle regulation is crucial for all living organisms and is often targeted by viruses to facilitate their own propagation, yet cell cycle progression control is largely underexplored in archaea. In this work, we reveal a cell cycle regulator (aCcr1) carrying a ribbon-helix-helix (RHH) domain and ubiquitous in the crenarchaeal order Sulfolobales and their viruses. Overexpression of several aCcr1 members including gp21 of rudivirus SIRV2 and its host homolog SiL_0190 of *S. islandicus* LAL14/1 results in impairment of cell division, evidenced by growth retardation, cell enlargement and an increase in cellular DNA content. Additionally, both gp21 and SiL_0190 can bind to the motif AGTATTA conserved in the promoter of several genes involved in cell division, DNA replication and cellular metabolism thereby repressing or inducing their transcription. Our results suggest that aCcr1 silences cell division and drives progression to the S-phase in Sulfolobales, a function exploited by viruses to facilitate viral propagation.

## INTRODUCTION

Viruses are dependent on host cellular machinery for propagation and therefore have evolved mechanisms to subvert host resources for their own benefit. One of the strategies is to inhibit cell division in order to keep host cells in a state favorable for DNA replication. Examples include many eukaryotic viruses (Fan et al., 2018), a couple of bacteriophages (Haeusser et al., 2014; Kiro et al., 2013; Stewart et al., 2013) and one family of archaeal viruses (Liu et al., 2021). While eukaryotic and bacterial viruses have been shown to inhibit cell division through direct protein-protein interaction with cell division machineries, how the archaeal virus STSV2 inhibit cell division is not known (Liu et al., 2021).

Archaeal cell cycle is reminiscent of that in eukaryotes displaying a gap 1 (G1) phase, a DNA synthesis phase (S phase), a long gap 2 (G2) phase and a rapid cell division (M and D) phase (Lundgren and Bernander, 2005). In addition, one of the archaeal large phyla, Crenachaeota, employs eukaryotic ESCRT (endosomal sorting complexes required for transport)-III and Vps4 proteins for cell division while another large phylum, Euryarchaeota, uses bacterial-like FtsZ for division (Caspi and Dekker, 2018). ESCRT-III and Vps4 coding genes have been identified in other archaeal genomes including those of many TACK (Thaumarchaeota, Aigarchaeota, Crenarchaeota and Korarchaeota) superphylum members and Asgardarchaeota (Caspi and Dekker, 2018), suggestive of similar cell division mechanisms in these organisms.

Several transcription factors are known to activate or inhibit transcription of genes needed for entering into, or exit from, the cell division phase in eukaryotes (Bähler, 2005; Galderisi et al., 2003) while much less is known about transcription factors controlling cell division in bacteria and archaea. A transcription factor conserved in many alphaproteobacteria, CtrA, controls cell division by binding to promoters of genes involved in cell cycle progression, as well as genes important for other cellular proceses (Francis et al., 2017). In archaea, the only known transcription factor controlling cell division is a ribbon-helix-helix (RHH) protein, CdrS, which activates transcription of division genes (Darnell et al., 2020; Liao et al., 2021).

RHH proteins are extremely widespread in Archaea, Bacteria and Eukarya. RHH domain containing proteins are DNA-binding and usually dimers. A previous study found RHH-containing proteins and glycosyltransferases to be extensively shared among crenarchaeal viruses, an outstanding feature given that viral families of the archaea domain share very few connector genes and hence appear to have distinct evolutionary histories (Iranzo et al., 2016). The distribution pattern of viral RHH transcriptional regulators suggested an important function during viral infection, and we set up to investigate the role of these proteins using rudivirus SIRV2 as our system of study. We demonstrate in this work that an RHH protein highly conserved in archaeal viruses and their host genomes inhibits cell division.

## RESULTS

### Homologs of the SIRV2 gp21 RHH-domain transcriptional regulator are widespread in crenarchaeal viruses

To identify the RHH-containing proteins in archaeal viruses, we annotated 153 archaeal viral genomes deposited in NCBI and retrieved all proteins harboring a RHH domain. This search recovered 194 genes from 90 genomes, mainly from crenarchaeal viruses and with only three highly divergent occurrences in euryarchaeal haloviruses (Table S1), similar to the taxonomic pattern observed by Iranzo et al. (Iranzo et al., 2016). Classification of the hits revealed that 115 of the 194 queries including SIRV2 gp21 (also known as ORF59b) belonged to the same cluster and that they are present in 87 viral genomes. The abundance and broad taxonomic distribution of the members of this cluster contrasts markedly with that of other viral RHH proteins, as the three largest clusters of the remaining queries contain only 19, 10 and 9 members each, including the previously studied transcriptional factor SvtR (Guillière et al., 2009), the homologs of SIRV4gp10, and the fuselloviral factor E73 (Schlenker et al., 2012). In addition, these clusters show a narrow distribution restricted to 1-3 viral families.

**Table 1.**
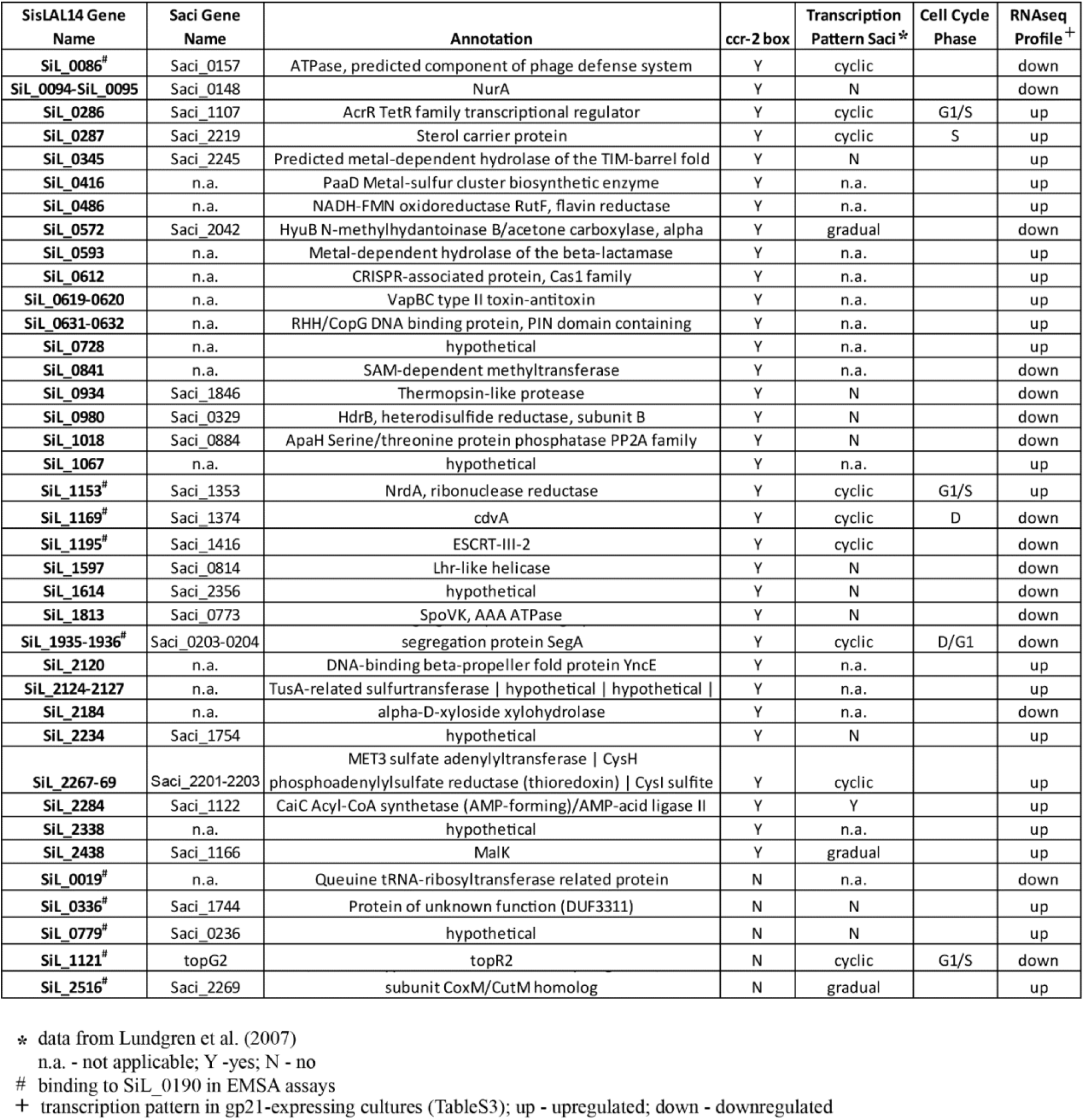
*S. islandicus* LAL14/1 promoters containing the ccr-2 box and transcription profile of their homologs in *S.acidocaldarius*.

| SisLAL14 Gene Name | Saci Gene Name | Annotation | ccr-2 box | Transcription Pattern Saci* | Cell Cycle Phase | RNAseq Profile<sup>+</sup> |
| --- | --- | --- | --- | --- | --- | --- |
| SiL_0086<sup>#</sup> | Saci_0157 | ATPase, predicted component of phage defense system | Y | cyclic |  | down |
| SiL_0094-SiL_0095 | Saci_0148 | NurA | Y | N |  | down |
| SiL_0286 | Saci_1107 | AcrR TetR family transcriptional regulator | Y | cyclic | G1/S | up |
| SiL_0287 | Saci_2219 | Sterol carrier protein | Y | cyclic | S | up |
| SiL_0345 | Saci_2245 | Predicted metal-dependent hydrolase of the TIM-barrel fold | Y | N |  | up |
| SiL_0416 | n.a. | PaaD Metal-sulfur cluster biosynthetic enzyme | Y | n.a. |  | up |
| SiL_0486 | n.a. | NADH-FMN oxidoreductase RutF, flavin reductase | Y | n.a. |  | up |
| SiL_0572 | Saci_2042 | HyaB N-methylhydantoinase B/acetone carboxylase, alpha | Y | gradual |  | down |
| SiL_0593 | n.a. | Metal-dependent hydrolase of the beta-lactamase | Y | n.a. |  | up |
| SiL_0612 | n.a. | CRISPR-associated protein, Cas1 family | Y | n.a. |  | up |
| SiL_0619-0620 | n.a. | VapBC type II toxin-antitoxin | Y | n.a. |  | up |
| SiL_0631-0632 | n.a. | RHH/CopG DNA binding protein, PIN domain containing | Y | n.a. |  | up |
| SiL_0728 | n.a. | hypothetical | Y | n.a. |  | up |
| SiL_0841 | n.a. | SAM-dependent methyltransferase | Y | n.a. |  | down |
| SiL_0934 | Saci_1846 | Thermopsin-like protease | Y | N |  | down |
| SiL_0980 | Saci_0329 | HdrB, heterodisulfide reductase, subunit B | Y | N |  | down |
| SiL_1018 | Saci_0884 | ApaH Serine/threonine protein phosphatase PP2A family | Y | N |  | down |
| SiL_1067 | n.a. | hypothetical | Y | n.a. |  | up |
| SiL_1153<sup>#</sup> | Saci_1353 | NrdA, ribonuclease reductase | Y | cyclic | G1/S | up |
| SiL_1169<sup>#</sup> | Saci_1374 | cdvA | Y | cyclic | D | down |
| SiL_1195<sup>#</sup> | Saci_1416 | ESCRT-III-2 | Y | cyclic |  | down |
| SiL_1597 | Saci_0814 | Lhr-like helicase | Y | N |  | down |
| SiL_1614 | Saci_2356 | hypothetical | Y | N |  | down |
| SiL_1813 | Saci_0773 | SpoVK, AAA ATPase | Y | N |  | down |
| SiL_1935-1936<sup>#</sup> | Saci_0203-0204 | segregation protein SegA | Y | cyclic | D/G1 | down |
| SiL_2120 | n.a. | DNA-binding beta-propeller fold protein YncE | Y | n.a. |  | up |
| SiL_2124-2127 | n.a. | TusA-related sulfurtransferase \| hypothetical \| hypothetical \| | Y | n.a. |  | up |
| SiL_2184 | n.a. | alpha-D-xyloside xylohydrolase | Y | n.a. |  | down |
| SiL_2234 | Saci_1754 | hypothetical | Y | N |  | up |
| SiL_2267-69 | Saci_2201-2203 | MET3 sulfate adenyltransferase \| CysH phosphoadenylsulfate reductase (thioredoxin) \| CysI sulfite | Y | cyclic |  | up |
| SiL_2284 | Saci_1122 | CaiC Acyl-CoA synthetase (AMP-forming)/AMP-acid ligase II | Y | Y |  | up |
| SiL_2338 | n.a. | hypothetical | Y | n.a. |  | up |
| SiL_2438 | Saci_1166 | MalK | Y | gradual |  | up |
| SiL_0019<sup>#</sup> | n.a. | Queuine tRNA-ribosyltransferase related protein | N | n.a. |  | down |
| SiL_0336<sup>#</sup> | Saci_1744 | Protein of unknown function (DUF3311) | N | N |  | up |
| SiL_0779<sup>#</sup> | Saci_0236 | hypothetical | N | N |  | up |
| SiL_1121<sup>#</sup> | topG2 | topR2 | N | cyclic | G1/S | down |
| SiL_2516<sup>#</sup> | Saci_2269 | subunit CoxM/CutM homolog | N | gradual |  | up |

\* data from Lundgren et al. (2007)
n.a. - not applicable; Y -yes; N - no

### The clade of SIRV2 gp21 homologs bears high similarity to a conserved cellular transcriptional regulator of Sulfolobales

The distribution pattern of RHH regulators in archaeal viruses suggests they were originally acquired from their hosts (Iranzo et al., 2016). A BLAST search to investigate the presence of cellular homologs of SIRV2 gp21 in the genome of *Sulfolobus islandicus* LAL14/1 revealed the presence of five small proteins with RHH-domains: SIL_RS00945 (also known as SiL_0190), SIL_RS07870 (SiL_1569), SIL_RS00595 (SiL_0120), SIL_RS14755 (no SiL_ annotation) and a homolog of SiRe_1373 (gene 1373 of *S. islandicus* REY15A) not annotated in the genome of LAL14/1 but located in the region between 1 286 819 to 1 286 682 bp. All of these correspond to highly conserved proteins in the Sulfolobales and possess identical genomic neighborhoods (Fig. S1A-E). Additionally, they appear to be essential to the host as revealed by a previous transposon mutagenesis and sequencing analysis of essential genes in *S. islandicus* (Zhang et al. 2018). A phylogenetic analysis of all viral and cellular homologs of SIRV2 gp21 splits them into several branches, five of which are represented each by one of the LAL14/1 homologs while four are composed uniquely of viral proteins. In addition, there are several interspersed viral homologs (Fig. 1). Interestingly, proteins from diverse viral families show h igh similarity to the clades of SiL_0190 and SiL_RS14755, with SIRV2 gp21 clustering with the former. While protein identity ranges between 100% −75.9% among the cellular homologs of SiL_0190, viral members of the clade share between 86.2% −26.1% identity to SiL_0190 (Fig. S2). We reasoned that such a high conservation within viruses that have little in common at the genomic level or mode of infection points out to a key function in their interaction with their crenarchaeal hosts.

**Figure 1.**
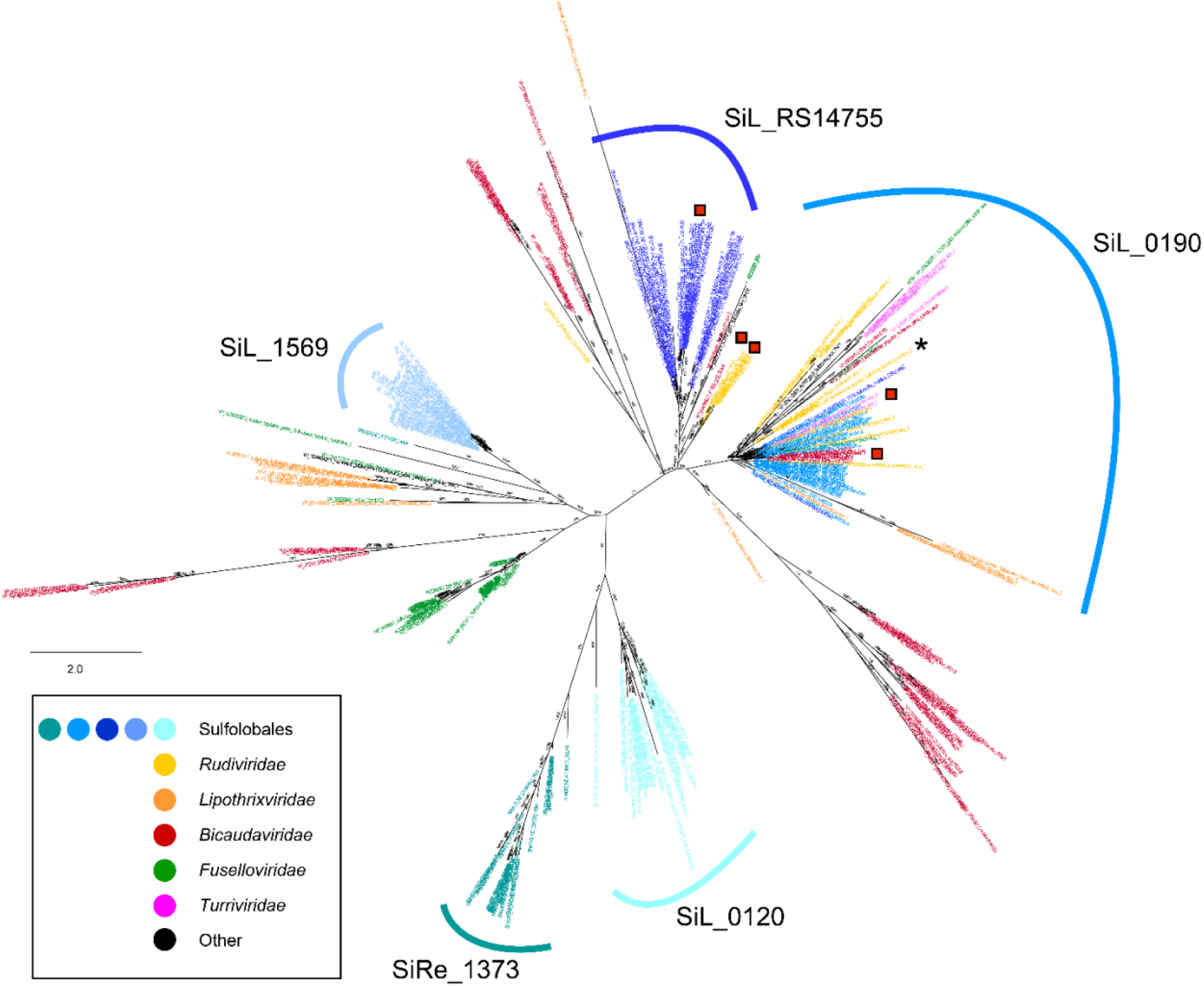
Phylogenetic analysis of the SIRV2 gp21 homologs in the Sulfolobales and their viruses. Color-coding is the same as in the inset panel with cellular homologs colored in shades of blue and viral homologs colored according to their viral taxonomy. The asterisk indicates the position of SIRV2 gp21 and the red squares indicate other transcriptional factors used in this work.

### SIRV2 gp21 regulates transcription of cellular genes involved in cell division and metabolism

SIRV2 gp21 is a 59-amino-acid protein (6.9 kDa) containing a ribbon-helix-helix motif indicative of transcriptional regulator activity (Schreiter and Drennan, 2007). It predicted to act as dimer with the N-terminal portion of the protein, which folds as a β-sheet, mediating site-specific interactions with the DNA (Fig. 2A). To investigate the role of SIRV2 gp21 during infection of its host *S. islandicus*, we first attempted to take advantage of the recently developed genetic engineering system for SIRV2 (Alfastsen et al., 2021; Mayo-Muñoz et al., 2018). However, our efforts to construct a viral strain encoding a His-tagged variant of gp21 or a knock-out mutant lacking gp21 were unsuccessful. Considering that gp21 is a conserved gene in all members of the rudiviridae and regarded as part of their core genome (Mayo-Muñoz et al., 2018), our results strongly suggest that gp21 is an essential gene for the virus. Initial efforts to identify the binding sequence of gp21 within the viral genome using recombinant protein expressed in, and purified from, *E. coli* showed nonspecific binding to DNA at increased protein concentrations (data no t shown). Therefore we cloned gp21 with a C-terminal His-tag into a plasmid vector for expression in *S. islandicus.* The tag was incorporated to the C-terminus which was shown to be unessential for the replication of SIRV2 (Mayo-Muñoz et al., 2018). Induction of gp21 expression triggered strong growth retardation (Fig. 2B), leading us to hypothesize that the target genes of this protein are located within the host genome. Chromatin immunoprotein sequencing (ChIP-seq) analysis to determine the binding sites of gp21 was, however, hindered by the low yield of the protein. Thus, we performed transcriptome profiling by RNA sequencing (RNA-seq) to compare gene expression remodeling after expression of gp21 (Table S2).

**Figure 2.**
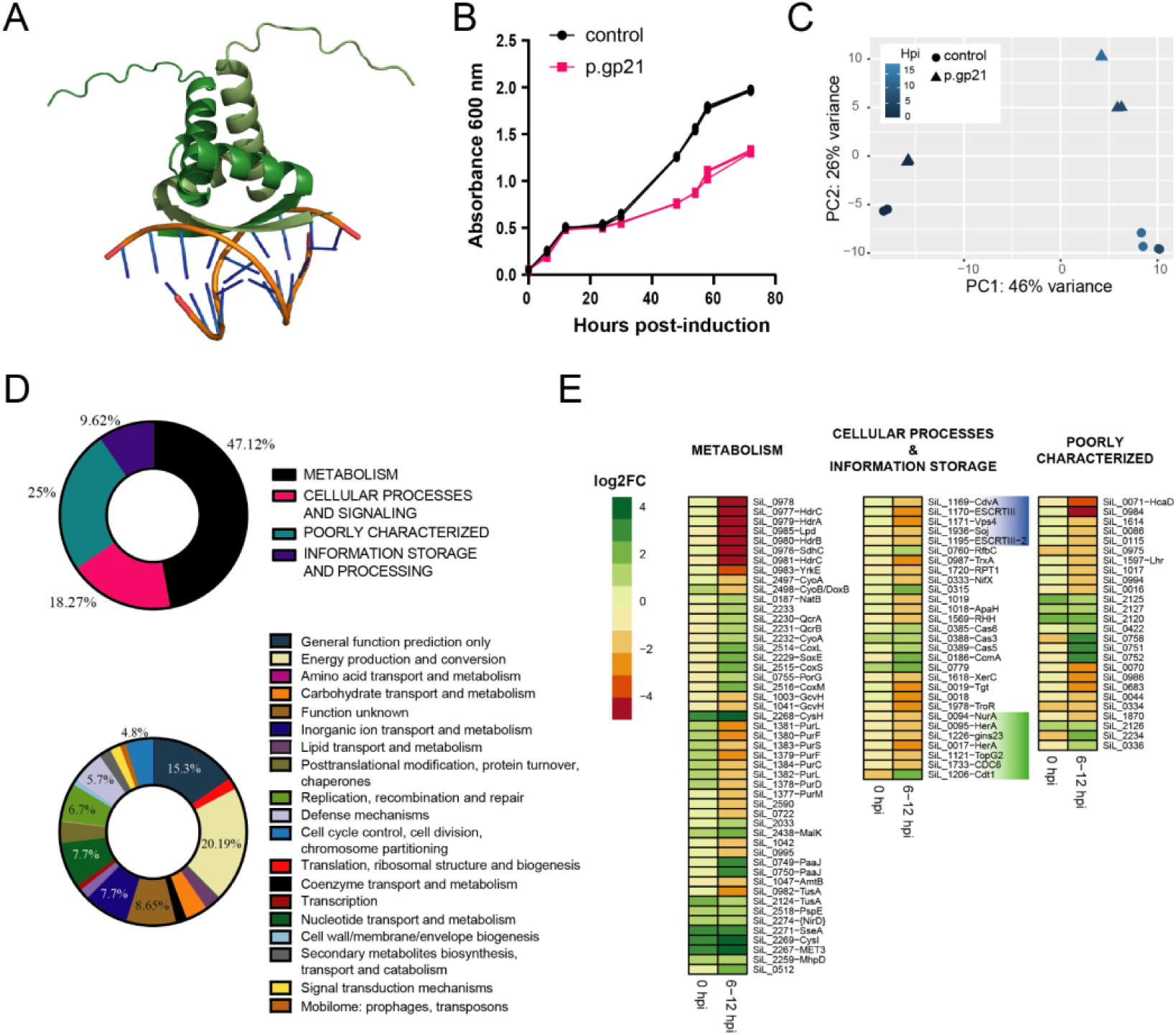
Expression of gp21 alters the transcription profile of genes involved in cell division and DNA replication. **A)** Model of the SIRV2 gp21 dimer. SIRV2 gp21 was modeled with AlphaFold2 and superimposed to the structure of the RHH-domain ArtA regulator with its target DNA (PDB: 3GXQ). Each chain of the gp21 dimer is indicated in different shades of green. **B)** Overexpression of gp21 elicits growth retardation. Growth in ACV medium of *S. islandicus* LAL14/1 harboring an empty plasmid (control) or a plasmid expressing gp21 under the control of the inducible arabinose promoter (p.gp21). Growth was monitored by measuring absorbance at 600 nm. Bars correspond to the standard deviation of three biological replicates. **C)** Principal component analysis of RNA-seq data from *S.islandicus* LAL14/1 cultures taken at 0, 6 or 12 hours post-induction (hpi) with arabinose. Control cultures harbor an empty plasmid, while p.gp21 indicates cells containing a plasmid for the expression of SIRV2 gp21. **D)** Clusters of Orthologous Proteins (COG) functional annotation of the genes selected as having a significant differential transcription in cultures expressing SIRV2 gp21 according to their COG-Module (top) or COG-Category (bottom) classification. E) Heatmap of the transcriptional profile of selected differentially expressed genes in p.gp21 vs. control cultures at 0 hpi with arabinose (basal gp21 expression) and at 6 and 12 hpi, colored by their log2 fold change and clustered by their COG-Module annotation. Genes belonging to the category of Cell cycle control, cell division and chromosome partitioning are shaded in blue. Genes of the category of Replication, recombination and repair are shaded in green.

The principal component analysis of the data shows that the control cultures (cells carrying the empty plasmid) group apart from cells expressing gp21 (Fig. 2C). By comparing control vs. gp21-expressing cultures i) before induction (at 0 hours post-induction) and ii) after induction, we observed 269 and 155 genes, respectively, that are differentially expressed in the gp21-expressing samples with a log2 fold-change over 1.4 (Table S3). We reasoned that the high number of differentially expressed genes could be the result of cascade effects initiated by the gp21-mediated changes and that if members of the SiL_0190 clade are so highly conserved, their target genes would also be conserved within the Sulfolobales, thus we used conservation as criterion to further filter the results. This allowed us to identify 103 genes differentially expressed, of which 8 showed strong modulation in the gp21-expressing cultures even before induction with arabinose, due to leaky expression of gp21 (Table S3, Fig. 2D). The majority of the differentially expressed genes belonged to the COG (Clusters of Orthologous Proteins) modules of Metabolism (47.6% of the genes) and Cellular processes and signaling (17.5%), followed by genes involved in Information storage and processing (9.7%) (Fig. 2E). A more detailed analysis of the functional categories revealed an overrepresentation of genes involved in Cell cycle control, division and chromosome partitioning (11-fold) and Replication, recombination and repair (2-fold) when compared to the distribution of COG terms in the genome of *S. islandicus* LAL14/1 (Fig. 2E). Among the metabolism-related categories, genes belonging to the Energy production and conversion (20% of the genes), and Inorganic transport and metabolism (7.7% of the genes) were preferentially regulated in our samples (Fig. 2E). Genes from the categories of Transcription, and Translation, ribosomal structure and biogenesis were depleted (5- and 6-fold, respectively) from our dataset.We were particularly interested in genes *sil_1169-sil_1171 (cdvA, escrt-III, vsp4), sil_1195 (escrt-III-2), sil_1935-1936 (segAB), sil_1121 (topG2), sil_1206 (cdt1), sil_1226 (gins23)*, and *sil_1733 (cdc6 1)*. Expression of the above genes was downregulated 2-6 times (1.5 −2.5 log2-fold) in the gp21-expressing cultures, with the exception of *sil_1206*, which was upregulated 2.8 times (about 1.7 log2-fold) (Fig. 2E, Supplementary Table S#). The first four are key components of the *Sulfolobus islandicus* cell division machinery and catalyze the last step in membrane abscission (Liu et al., 2017). The operon of *sil_1935-36* code for the components of the segrosom SegAB, involved in chromosome organization prior to partitioning (Yen et al., 2021). TopG2 is a reverse gyrase presumably involved in DNA replication and repair in the crenarchaea (Couturier et al., 2020). Cdt1 and Cdc6-1 recognize the origins of chromosome replication, and Gins23 is part of the MCM complex, the motor driving unwinding of DNA at the replication fork (Lang and Huang, 2015; Samson et al., 2013). Among the genes assigned to the Metabolism category, components of the heterodisulfide reductase complex (Hdr) were strongly downregulated, as well as genes involved in purine biosynthesis (*sil_1377-1384*) (4 and 2 log2-fold, respectively). On the other hand, genes coding for cytochrome subunits (sil_2229-2233) were upregulated (1.5 − 2.3 log2- fold), similarly to several genes in the Inorganic ion transport and metabolism (e.g. *sil_2267-2271*) (3.3 - 4.4 log2-fold. These results led us to speculate that gp21 is involved in the regulation of genes required for cell division and genome replication.

### Overexpression of members of the Sil_0190 clade triggers growth arrest and cell enlargement

We reasoned that inhibition of cell division genes in gp21 overexpressing cells could be related to the growth arrest phenotype observed in these cultures. We performed microscopy and flow cytometry analyses of cells at different time-points after induction of plasmid-borne gp21 or an empty plasmid as a control. Surprisingly, we observed a remarkable increase in cell size noticeable after 24 hours post-induction (hpi) (Fig. 3A), when 66.7 ± 2.3% of the cells showed cell enlargement equivalent to a 35 fold increase in the cell volume (Fig. 3B). Cell enlargement continued with incubation time and cells at 72 hp i had expanded their diameter up to 5-fold, corresponding to an increase in volume of approximately 150 times (Fig. 3B). Additionally, we observed an increment in the number of genome equivalents within the population of large cells. This increment was significant already by 12 hpi when cells mostly contain over two chromosome equivalents (2C) and peaked after 30 hpi, with the majority of the cells containing between 4-10 genome equivalents (Fig. 3A). Yet, we observed a fraction of the population showing th e characteristics of normal cell growth, with respect to size and genome content, that increased from 33% at 24 hpi to 45% at 72 hpi (Fig. 3A). This fraction of cells probably explains the continued, albeit slower, growth observed for the gp21-expressing samples (Fig. 2A).

**Figure 3.**
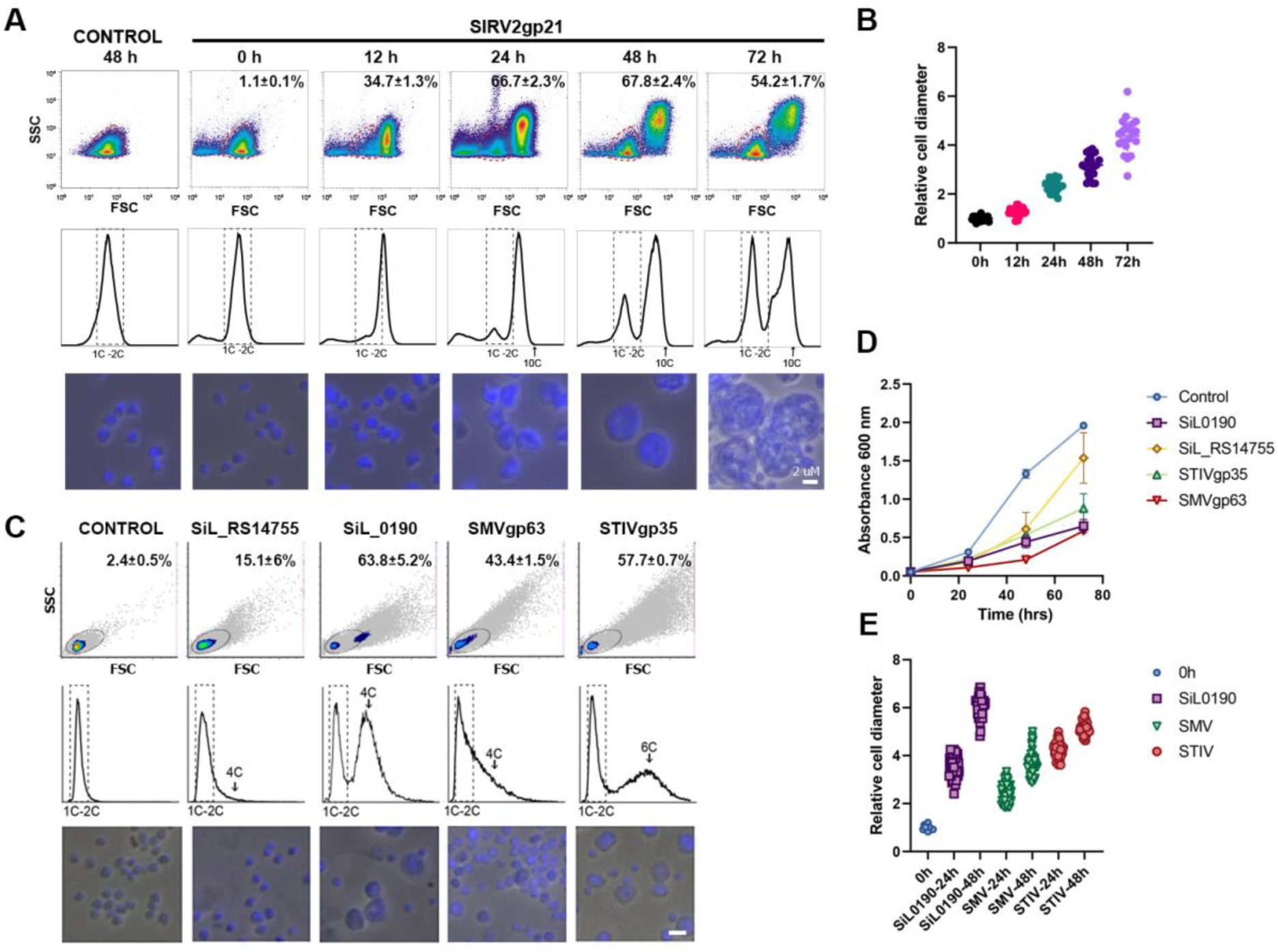
Members of the SiL_0190 clade induce cell enlargement and increased DNA content. **A)** Effects of expression of SIRV2 gp21 on cell size and DNA content at different time-points after induction measured by flow cytometry (top and middle panels) and microscopy (bottom panel). Bar = 2 um. The control corresponds to cells harboring an empty plasmid at 72 hpi. The percentage of large cells is indicated in the graphs of cell size. The square indicates cells with normal DNA content corresponding to 1-2 chromosome copies**. B)** Average cell diameter of 25 gp21-expressing cells. Bars correspond to the standard deviation of the data. **C)** Distribution of cell size and DNA content (top and middle panels) and microscopy images (bottom panel) of cultures expressing viral and cellular homologs of the SiL_0190 clade, an empty plasmid (control) or SiL_RS14755 at 24 hpi. **D)**Growth curves after induction with arabinose of *S. islandicus* strains harboring an empty plasmid (control), or plasmids expressing SiL_RS14755, SiL_0190, STIV gp25 or SMV gp63. The graph shows the average and the standard deviation of three biological replicates. **E)** Average cell diameter of 50 random cells from the cultures in panel C at 24 or 48 hours post-induction of protein expression.

The enlarged-cell phenotype with cells containing more than two genome equivalents has been previously observed in *Sulfolobus* after silencing of cell division genes, in cells overexpressing dominant-negative mutants of the Cdv complex and in cells where cell division has been inhibited as a result of virus infection (Liu et al., 2017, 2021; Samson et al., 2008, 2011; Yang and Driessen, 2015), suggesting that overexpression of SIRV2 gp21 results in inhibition of cell division, but not DNA replication. Moreover, the high conservation of members of the clade among different families of the Sulfolobales and crenarchaeal viruses suggests that the homologs perform an equivalent function. To explore the functional equivalence of the members of this branch, we transformed a plasmid with an inducible-expression cassette containing RHH-domain proteins of the SiL_0190 and SiL_RS14755 phylogenetic branches, namely the cellular SiL_0190 and SiL_RS14755 and the viral SIFV gp29 from a lipothrixvirus, the turriviral STIV gp35, SMV3 gp63 from a bicaudavirus and the rudiviral SIRV9 gp19, or an empty vector as a control. All these proteins are predicted to fold into an RHH motif and to dimerize, with the exception of SIFV gp29, which already contains two RHH motifs in the same peptide chain and folds as a monomer (Supplementary Figure 3). We subsequently monitored changes in cell growth curve, size and genome content occurring after induction of target gene expression (Fig. 3C). All the constructs transformed with the exception of the vector harboring SiL_RS14755, showed a 10-fold reduction in their transformation efficiency when compared to the empty vector and a small colony phenotype suggestive of slower growth, and induced growth retardation after induction of protein expression (Fig. 3D), similar to what was observed with SIRV2 gp21. We were unable to grow the strains harboring a plasmid for SIFV gp29 expression in liquid culture, implying that leaky expression of these factors already causes aberrant growth.Overexpression of SiL_0190 triggered similar changes as we observed for gp21: 63.8 ± 5.2% of the cells increased their size by 24 hpi, decreasing slightly to around 40% with prolonged incubation, while the diameter of this fraction of the population increased up to around 6-times by 72 hpi equivalent to a raise in cell volume of 150-fold (Fig. 3E). Moreover, the enlarged cells contained over 6 genome equivalents after 72 h of incubation (Fig. 3C). The viral homologs STIV gp35 and SMV3 gp63 also triggered cell enlargement and increased DNA content, albeit in a milder way and with the strongest phenotype seen at 24 hpi, when 57.7 ± 0.7% and 43.4 ± 1.5% of the cells had increased their diameter by 4- and 3-fold, respectively (Fig. 3F). Contrastingly, the fraction of the population corresponding to large cells decreased during incubation, stabilizing at 20-25%. Whereas some STIV gp35-expressing cells contained around 6 genome equivalents, cells harboring SMV3 gp63 showed a moderate doubling of their DNA content equivalent to 3-4 chromosome copies (Fig. 3F). The milder phenotype caused by the turriviral and bicaudaviral fractions could reflect differences between the native hosts of these viruses and *S. islandicus* LAL14/1. On the other hand, overexpression of SiL_RS14755 did not affect cell size or genome content, indicative of a different function of the members of this clade. Overall, our results support a conserved role of the SiL_0190 clade in the regulation of cell division in the Sulfolobales.

To further investigate if SIRV2 gp21 and SiL_0190 were functionally equivalent, we attempted to substitute *gp21* in the genome of SIRV2 with *siL_0190* while keeping intact the native regulatory regions of the former. We employed a CRISPR-based genome editing strategy where a strain harboring a plasmid containing a spacer targeting *gp21* and a donor DNA fragment consisting of SiL_0190 flanked by the flanking sequences of *gp21* in the viral genome was used to facilitate recombination after infection (Alfastsen et al., 2021). We were unable to obtain the recombinant virus after several attempts, suggesting that SiL_0190 cannot complement the function of SIRV2 gp21 during viral infection.

### Members of the SiL_0190 clade of transcriptional regulators bind to a sequence motif AGTATTA

To determine the mechanism by which SiL_0190 homologs inhibit cell division in Sulfolobales, we investigated possible binding of SiL_0190 and SIRV2 gp21 to the promoter region of the differentially expressed genes identified in the RNA-seq experiment and to viral promoters, including the promoter of *gp21*. For this, we performed competition assays where a recombinant protein was mixed with 200 bp dsDNA fragments corresponding to the labeled promoters of the target genes and different concentrations of competitor DNA. The results of PAGE-EMSA reveal three patterns: i) no DNA binding, as evidenced by a lack of retardation in DNA migrati on after electrophoresis (e.g. SIRV2 *gp01* promoter, Fig. 4A-B top row), ii) unspecific, low affinity binding, quickly displaced by the addition of competitor DNA in a ratio of 1:10 (e.g. the promoters of the bacteriophage T7 polymerase, SIRV2 *gp15*, or *sil_0985,* Fig. 4A-B top row), and iii) specific binding evidenced by retardation of DNA migration at high concentrations of competitor DNA up to 1:50 (Fig. 4A-B). gp21 and SiL_0190 bind fifteen and eleven DNA probes, respectively, in a specific manner. Promoters bound by both proteins include the regulons of the cell division machinery components *sil_1169* (*cdvA)* and *sil_1195* (*escrt-III-2*); the regulatory sequences of genes involved in DNA metabolism *sil_1121* (*topG2*)*, sil_1936* (*segA*) and the promoters of *sil_0086* (uncharacterized AAA-type ATPase), *sil_0336* (DUF3311), *sil_0779* (hypothetical protein), *sil_1153* (NrdA, ribonucleotide reductase) and *sil_2516* (CoxM, glyceraldehyde oxidoreductase subunit). In addition, gp21, but not SiL_0190, specifically recognizes the promoters of *sil_1206* (Cdt1), *sil_0095* (HerA-like helicase) and *sil_0979* (hypothetical protein), plus the viral promoters of the middle-late genes *gp17*, *gp20-gp21* (Fig. 4A-B, bottom row). These slight differences in target preference suggest that gp21 may have additional functions than the host factor that explain why SiL_0190 was unable to substitute for gp21 (data not shown).

**Figure 4.**
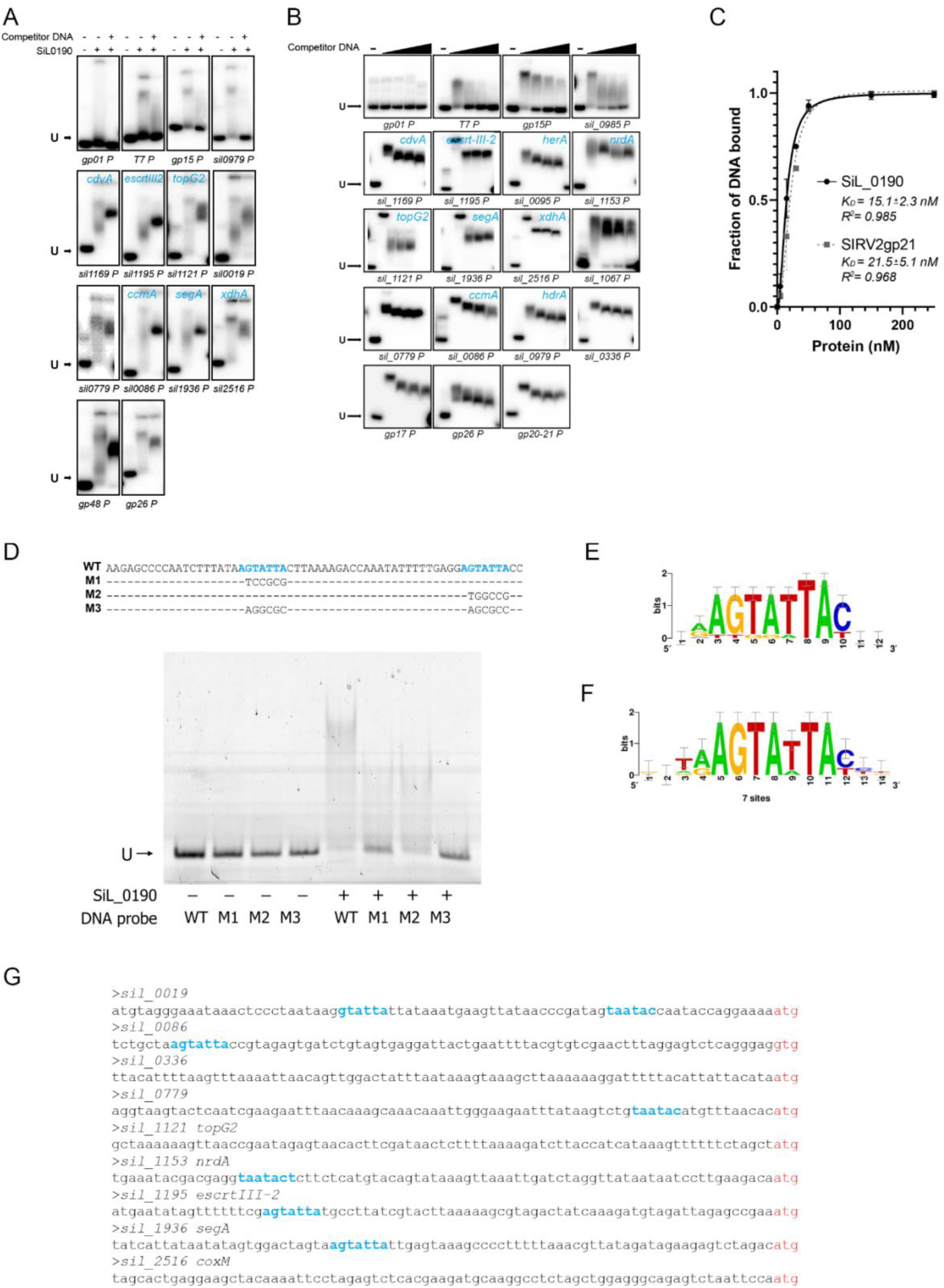
SiL_0190 binds to the motif in the promoters of cell division and DNA replication genes. **A)** Representative results of mobility shift assays for SiL_0190. A total of 30 ng labeled probe was mixed with increasing concentrations of cold competitor DNA (ratios of 1:1, 1:10) and 100 nM protein. The first lane of every experiment, marked with a “-”, corresponds to the labeled probe without protein nor competitor DNA. The name of the probe is indicated below each experiment. Arrows point to the unbound (U) probe. **B)** Representative results of mobility shift assays for SIRV2 gp21. Assays were conducted as described in panel A with additional concentrations of cold competitor DNA (ratios 1:1, 1:10, 1:20 and 1:50). **C)** Apparent binding affinities (KD) of SiL_0190 and SIRV2 gp21. Curves represent the average of shifted probe as a function of protein concentration. Average data from three independent experiments, bars indicate the standard deviation. **D)** Top: Upstream region of the *cdvA* gene of *S. islandicus* LAL14/1 and variants with mutations of the AGTATTA motif (highlighted in blue). Bottom: EMSA assay results for the *cdvA* promoter variants in panel D for SiL_0190. **E-F)** Enriched motifs identified in the SiL_0190-**(E)** or the SIRV2gp21-bound promoters **(F). G)** Occurrences of the DNA-binding motif AGTATTA in the target genes of SiL_0190.

We determined the dissociation constant (KD) of the interaction between SiL_0190 and gp21 to the *cdvA* promoter (*sil_1169*), 15.1 nM and 21.5 nM, respectively (Fig. 4C), showing that the affinity to DNA of the two proteins is very similar. Binding to the promoter of *cdvA* is also highly specific, as the dissociation constant of SiL_0190 remains the same in the presence of excess competitor DNA (Supplementary Figure S4). To determine the binding site of SiL_0190 we used 100 bp sequences of the target promoters to predict enriched motifs with the RSAT oligo-analysis tool (van Helden et al., 1998; Nguyen et al., 2018). The analysis identified two motifs enriched in the SiL_0190-bound promoters with the seed sequence GAGTCT and AGTATTA (Supplementary Figure 5A-B). Mutagenesis along the promoter of *cdvA* abolished binding only when the motif AGTATTA was absent, suggesting this oligonucleotide represents the SiL_0190 binding site (Supplementary Figure 6). This motif is present twice in the *cdvA*-regulatory region, between positions −10 to −2 and −40 to −46 (Figure 4D), hence we specifically mutated each copy and evaluated its contribution to the binding of both SiL_0190 and SIRV2 gp21 (Figure 4D-E). We observed impaired binding when only one copy was present, and, mutation of both copies was necessary to completely abrogate the binding (Figure 4E, Supplementary Figure 6C), demonstrating that AGTATTA is the DNA-binding motif of SiL_0190 and gp21. We additionally tested binding to the promoter of *sil_1936* that has a single occurrence of the motif, observing abrogation of DNA binding capacity after its mutation (Supplementary Figure 6D-E). The SiL_0190-binding motif is present, however, in only five of eleven bound targets (Figure 4A, 4F), thus it remains to be determined which factors mediate recognition of the other promoters. On the other hand, we were able to identify the binding-motif in the 150 bp upstream region of 240 genes of *S. islandicus*, of which 33 were differentially expressed in the RNAseq experiment (Table 1).

We performed quantitative RT-qPCR to investigate whether SiL_0190 regulates expression of genes that were up- or down-regulated by SIRV2 gp21 (Figure 2). RNA was extracted from cells overexpressing Sil_0190 and the subsequent RT-qPCR demonstrated gene regulation highly similar to what was observed for *gp21*-overexpressing cells (Supplementary Figure S7). Among the tested genes, *cdvA*, *escrtIII-2* and *segA* were downregulated −2.5, −3.2 and −2.8 log2 fold and the ribonucleotide reductase genes were upregulated 2.2 log2 fold by Sil_0190 overexpression.

To investigate if the other members of the clade share the same DNA-binding motif, we predicted enriched oligonucleotides in the promoters of the homologs of the SiL_0190 target genes in four different members of Sulfolobales (Supplementary Figure 5). A motif with the seed sequence GTATTA was identified in three of the four species and occurrences of the seed were identified in the analyzed promoters of the four organisms, in particular the promoters of *cdvA*, *escrtIII-2* and *nrdA* consistently contain the motif in all cases analyzed (Supplementary Figure 8).

Earlier transcriptome analyses showed that SIRV2 *gp21* expression was detectable at 30 min post-infection and reached maximum at 1 h (Okutan et al., 2013; Quax et al., 2013), and the host gp21-targets identified here followed the same regulation pattern (repression or activation) (Supplementary Figure S9). For example, *cdvA* was found to be downregulated around 2.2 log2 fold at 2 hours post SIRV2 infection and the ribonucleotide reductase genes were shown to be upregulated 2.1 log2 fold at the same time point. This strongly suggests that these changes were due to the activity of gp21 and that the gene regulation following gp21 overexpression observed in this study is biologically relevant.

### SiL_0190 is a cell cycle regulator that drives the transition between the D- and S- phases of the cell cycle

Previous work on *Sulfolobus acidocaldarius* identified *saci_0942* (the homolog of SiL_0190 in this organism) as a putative regulator of the transition to the stationary phase (Lundgren and Bernander, 2007). Additionally, Lundgren et al. (Lundgren and Bernander, 2007) identified a subset of genes sharing a similar transcription pattern and harboring the consensus element TGTATTAT, which they named as ccr-2 (cell cycle regulon 2) box. This motif resembles the SiL_0190 binding motif identified in this work (AGTATTA) and the latter is present in the promoters of the homologs of the ccr-2 box-containing genes in *S. islandicus*. We therefore name the gp21/SiL_0190 clade as aCcr1 (archaeal <u>c</u>ell <u>c</u>ycle regulator <u>1</u>), the first master regulator of the cell cycle characterized in Sulfolobales.

It was reported that induction of saci_0942 occurred during the G1/S transition (Lundgren and Bernander, 2007). Interestingly, the genes regulated by aCcr1 (gp21/SiL_0190) overexpression in *S. islandicus* LAL14/1 exhibited a consensus expression pattern in the synchronized *S. acidocaldarius* if a respective homolog is present in the latter genome and the time-course expression data was described (Lundgren and Bernander, 2007). That is, those downregulated by aCcr1 shown in this work (*cdvA, escrtIII-2, segA, topR2*) were induced in an earlier stage (D- or D/G1) than a*ccr1* whereas those upregulated by aCcr1 (*saci_1107, saci_2209*, see Table 1) were induced at a later stage in the cell cycle (S-phase) (Figure 5, Table 1). This strongly suggests that aCcr1 homologs regulate cell cycle progression in Sulfolobales by inhibiting the transcription of cell division genes and other D/G1-phase genes, and by inducing transcription of S/G2-phase genes.

**Figure 5.**
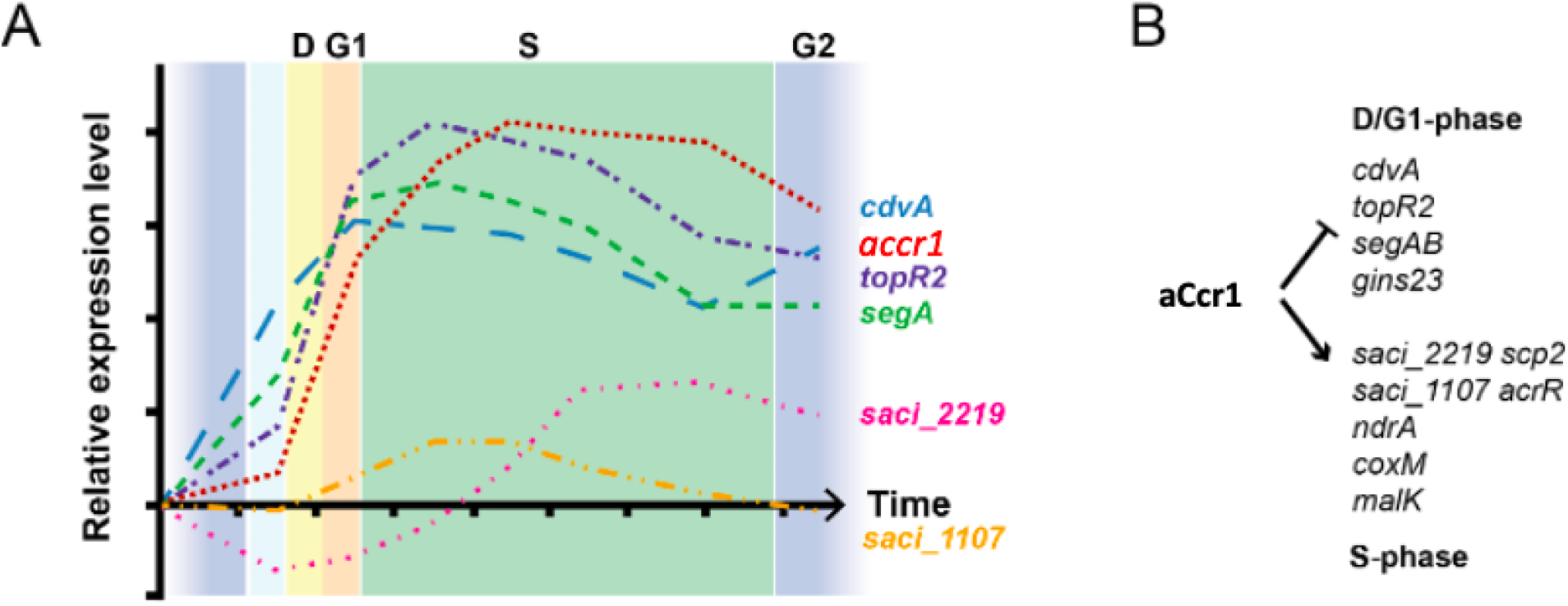
Transcriptional profile of the homologs of some aCcr1 targets in *S. acidocaldarius*. **A)** Expression profiles of cyclic genes in *S. acidocaldarius* as determined by Lundgren et al. (2007). The homologs in *S. islandicus*LAL14/1 of the genes shown contain the ccr-2 box in their promoter region and are recognized by SiL_0190 and SIRV2 gp21. **B)** Effect of the overexpression of aCcr1 in *S. islandicus* over the transcription of the corresponding homologs of the genes shown in panel A. Induction of aCcr1 correlates with the decrease or plateau of genes induced earlier during the D- and G1 phases (e.g. *cdvA, segA, topR2, gins23*) and with the induction of S-phase genes (e.g. *scp2, acrR*).

## DISCUSSION

The pioneer work by Lundgren et al. first described cell-cycle specific transcription of over 160 genes in the crenarchaeaon *Sulfolobus acidocaldarius* (Lundgren and Bernander, 2007), pointing to transcriptional regulation as a mechanism for the control of cell cycle progression in Sulfolobales. However, no cell-cycle regulatory factor has been described so far for Crenarchaeaota. In this work, we identify and characterize aCcr1, a conserved, essential protein in Sulfolobales that functions as a regulator of cell cycle progression. aCcr1 downregulates the transcription of important components of the Cdv (cell division) machinery CdvA and CdvB2, SegA, a component of the chromosome segregation system (Kalliomaa-Sanford et al., 2012) and the reverse gyrase TopR2 that regulates DNA supercoiling in crenarchaeota (Garnier et al., 2021). It additionally upregulates the transcription of the ribonucleotide reductase NrdA that converts ribonucleotides into deoxyribonucleotides, an essential step for DNA synthesis, and CoxM, a component of the CO dehydrogenase important for CO oxidation in Sulfolobus (Sokolova et al., 2017). aCcr1 binds to the motif AGTATTA (ccr2-box, Fig. 4) that is present in the promoter region of several of its target genes. aCcr1 is highly conserved in Sulfolobales, with amino acid identities between 81-100% (Supplementary Figure 2A) similarly to its target genes, which all have homologs in all Sulfolobales and whose promoters also contain the the motif AGTATTA (Supplementary Figure 7).

Our knowledge about cell cycle control in archaea is still very limited. The cell cycle transcriptional regulator CdrS (cell division regulator short) present in the Euryarchaeota phylum, was recently identified and characterized (Darnell et al., 2020; Liao et al., 2021). CdrS harbors an RHH domain that binds to promoters, and activates the transcription, of 18 genes, including four involved in cell division (*ftsZ1, ftsZ2*), DNA replication and partitioning (*sepF* and *smc*) and several genes involved in metabolism (Darnell et al., 2020; Liao et al., 2021). While only two cell cycle regulators have been described in the archaea domain (CdrS and aCcr1), it is interesting to note that both contain the RHH structural motif, whereas the majority of the transcriptional factors encoded in archaeal genomes harbor helix-turn-helix (HTH) motifs (Flores-Bautista et al., 2020; Lemmens et al., 2019). On the other hand, RHH transcriptional factors are the most abundant in archaeal viruses (Iranzo et al., 2016). The common use of RHH transcriptional factors despite the fact that cell division machineries between the Euryarchaeota and the Crenarchaeota are completely different (the tubulin FtsZ system and the ESCRT-based machinery, respectively, reviewed by (Caspi and Dekker, 2018; Ithurbide et al., 2022)) suggests a common theme in cell cycle control in the Archaea domain. RHH-domain proteins have so far been described solely in bacteria and archaea (Schreiter and Drennan, 2007). Intriguingly, while transcriptional control appears as common theme in cell cycle regulation for both bacteria and eukaryotes, to our knowledge no transcriptional factor of the RHH family has been implicated in cell cycle control in bacteria. Additional work is required to elucidate the role of RHH-domain transcriptional factors in regulating cell cycle progression in the Archaea domain.

We predict the existence of several regulatory factors whose successive activation drives the transcription of different sets of genes during cell cycle progression. A similar mechanism has been described in the alphaproteobacteria, where a circuit of antagonistic regulators act together to control transcription across the different phases of the cell cycle (reviewed Panis et al. 2015). In eukaryotes, transcriptional control of the cell cycle is linked to the activity of cyclins and cyclin dependent kinases, which, among other changes, regulate the activity of the three RNA polymerases (Delgado-Román and Muñoz-Centeno, 2021).

Regulation of cell growth, cytokinesis and DNA replication and segregation is exquisitely and robustly coupled in the three domains of life. These processes are linked together by several layers of regulation that involve different mechanisms and control strategies, and we envisage a similarly complex picture for the archaea domain. For example, post-translational control of the cell division machinery by the proteasome was reported in *Sulfolobus acidocaldarius*, where the 20S proteasome degrades CdvB and triggers cell division by allowing the constriction of the CdvB1:CdvB2 division ring (Tarrason Risa et al., 2020). Post-translational mechanisms could be involved in regulation of aCcr1, since plasmid-driven overexpression of aCcr1 homologs in this work constantly resulted in low protein yield, despite strong transcription from the arabinose promoter.

The widespread presence of aCcr1 homologs in crenarchaeal viruses is striking, as it encompasses six different families of viruses that have diverse hosts and infection modes, including both lytic and lysogenic viruses (Figure 1). This highlights the importance of this protein for successful viral infection. The pressure to maintain sequence and functional conservation of aCcr1 homologs is remarkable, as exemplified by the high identity (over 40%) at the amino acid level of the viral aCcr1 homologs and the fact that no other protein is shared among so many archaeal viral taxons. We hypothesize that SIRV2 and other viruses benefit from regulating cell cycle by driving the host cell into a state where components important for viral replication are present. For example, the ribonucleotide reductase NdrA (*sil_1153*), one of the highly activated genes following SIRV2 infection, has been proposed to be important for viral replication by converting ribonucleotides into deoxyribonucleotides for DNA replication and repair (Okutan et al., 2013).

There are multiple reports of eukaryotic viruses regulating cell cycle during infection through different mechanisms (reviewed by (Panis et al., 2015)), and a few bacterial viruses that mostly disrupt proper functioning of the cell division protein FtsZ (Bhambhani et al., 2020; Faubladier and Bouché, 1994; Hernández-Rocamora et al., 2015; Kiro et al., 2013; Ragunathan and Vanderpool, 2019). There is only one report of an archaeal virus family putatively disrupting the cell cycle of its host, infection with viruses of the *Bicaudaviridae* family STSV2 and SMV1 arrests cells in the S-phase and triggers transcriptional repression of components of the ESCRT cell division machinery, resulting in cell enlargement and an increase in the number of chromosomes per cell (Liu et al., 2021). Interestingly, some bicaudaviruses, namely ATV, ATV2 and SMV3, encode homologs of aCcr1 while STSV2 contains several members of the gp21 family of transcriptional regulators, none of these belong to the aCcr1clade, hence it remains to be determined if the RHH-domain factors in STSV2 are involved in cell cycle control.

The family of RHH-domain regulators to which aCcr1 belongs contains many branches that include several cellular and viral proteins. In this work, we have functionally characterized the clade of aCcr1, whose special feature is the taxonomic diversity of its members, as a transcriptional regulator that drives progression into the S-phase of the cell cycle. Our preliminary data suggests that other branches of the family may perform novel, different functions in Sulfolobales, and future work will shed light on the role of these cellular and viral clades of the family.

## METHODS

### Strains and growth conditions

Cultures of *Sulfolobus islandicus* LAL14/1 Δarrays (He et al., 2016) were grown at 78°C in SCVU medium (basic salts medium (Zillig et al., 1993) supplemented with 0.2% sucrose, 0.2% casamino acids, vitamin mixture and 0.002% uracil) with agitation. After plasmid complementation of the ΔpyrEF mutation, uracil was removed from the medium. Induction of protein expression and knockdown cassettes was done by growing cells in ACV medium, which has the same composition as SCV but substitutes sucrose for 0.2% D-arabinose. *Escherichia coli* strains DH5a and Rossetta (DE3) were used for plasmid cloning and protein expression, respectively, and grown in LB medium supplemented with the appropriate antibiotics.

### Virus propagation and infection of cell cultures

*S. islandicus* LAL14/1 cultures were grown until the optical density (OD) at 600nm is 0.2, then infected with SIRV2 and incubated for around 48h until the OD_600_ decreased. Cell were removed by centrifugation at 5000 x g for 10 min and the virions in the supernatant were recovered by addition of 10% polyethylene glycol 6000 (PEG6000) and 1M NaCl, followed by an overnight incubation at 4℃. Virus by collected by centrifugation at 11000 x g for 30min and resuspended in 10 mM Tris-acetate pH 6. Remaining cell debris was removed by centrifugation at 11000 x g for 5min. Virus titer was determined by plaque assay as described previously (Alfastsen et al., 2021). Briefly, 100 uL of ten-fold serial dilutions of the virus were added to 2mL fresh *S.islandicus* LAL14/1 △array cells (OD_600_ around 0.2) and incubated at 78℃ for 30min. Then, the infected cells were mixed with an equal volume of 0.4% Gelzan and spread onto the pre-warmed 0.7% Gelzan/SCV plates. Plates were incubated at 78℃ for 48 h and plaques formed were counted to determine the virus titer.

### Plasmids, oligonucleotides and DNA sequencing

A list of the plasmids and oligonucleotides used in this study is provided in Tables S5-S8. Oligonucleotides and gBlocks were synthesized by Integrated DNA Technologies, Inc. All constructs were verified through Sanger sequencing, using the services of Eurofins Genomics.

### Plasmid construction

Plasmids were constructed using the vectors pGE2 and pEXA2 as backbones. The plasmid pEXA2 is a shuttle vector, it contains a *pyrEF* cassette as a selective marker in *Sulfolobus* (Peng et al., 2012) and pGE2 is a derivative of pEXA2 (He et al., 2016). Vector pGE2-MCS was constructed by inserting multiple cloning site 1 (MCS1) synthesized as a gBlock into pGE2 upstream of the pEXA_Sy primer binding-site using overlapping-end PCR.

SIRV2 gp21 was amplified from SIRV2 genomic DNA (NC_004086.1), the DNA fragment obtained was inserted between the NdeI and NotI restriction sites of pGE2, resulting in Sis/pGE2- SIRV2gp21(Supplementary Table S5). The PCR products were later purified with the PCR purification kit (GeneJET K0702, Thermo Scientific).

The viral homologs of the protein SIRV2gp21 were selected from lipothrixvirus (SIFV0029 NP_445694.1), turrivirus (STIV-A61 YP_024996.1), bicaudavirus (SMV3gp63 YP_009226286.1), and usarudivirus (SIRV9gp19 YP_009362571.1). In addition, the archaeal factors SiL_0190 and SiL_RS14755 were amplified from the host *Sulfolobus islandicus* LAL14/1 Δarrays (He et al., 2016), whereas gBlocks were synthesized for the viral factors SIFV0029, STIV-A61, SMV3gp63 and SIRV9gp19 (Supplementary Table S5).

### Construction of a gp21-knockout viral strain

Plasmids carrying CRISPR spacers and homologous sequences for recombination were constructed using protocols described earlier (Li et al., 2016; Mayo-Muñoz et al., 2018). Briefly, spacer fragments were generated by annealing of the corresponding complementary oligonucleotides and inserted in the LguI restriction sites of pGE2-MCS to yield plasmids carrying an artificial mini-CRISPR array; then, donor DNA fragments for homologous recombination were obtained by extension overlap PCR and inserted between the NheI and AvrII restriction sites (Supplementary Table S6). The resulting plasmid *Sis*/pGE2-d-SIRV2 gp21 was transformed by electroporation into *S. islandicus* LAL14/1 Δarrays as described previously (Deng et al., 2009).

### Protein expression and purification

The genes SIRV2 gp21 and SiL_0190 were PCR-amplified from the SIRV2 and *S.islandicus* LAL14/1 genomes, respectively, and inserted into the expression plasmid pET30a (Novagen). Recombinant SIRV2 gp21 and SiL_0190 with C-terminal His-tag were expressed using *Escherichia coli* Rosetta^TM^ (DE3) cells (Novagen). Protein expression was induced during logarithmic growth through the addition of 0.5mM isopropyl-thiogalactopyranoside (IPTG) followed by an overnight incubation at 16℃. Cells were collected by centrifugation and the cell pellet was resuspended in lysis buffer (50 mM HEPES pH 7.4, 300 mM NaCl, 20 mM imidazole). Cells were lysed by sonication and French press and the cell extract was clarified by centrifugation at 13000 g for 30 min at 4 ℃. The supernatant was loaded onto a 5mL His-trap column (GE Healthcare) for protein purification and eluted with a linear imidazole (20 mM-200 mM imidazole) gradient in lysis buffer. Fractions were pooled and concentrated to 1mL and purified further by the size exclusion chromatography (SEC) using a Superdex 200 column using the chromatography buffer (50 mM HEPES pH 7.4, 300 mM NaCl).

### RNA sequencing

Samples collected at 0, 6, 12 and 24 hours post-induction with arabinose from duplicate cultures of *S. islandicus* LAL14/1 Δarrays harboring an empty pGE2 plasmid (control) or a plasmid expressing SIRV2 gp21. Total RNA was extracted using TRI reagent (Sigma-Aldrich) and RNA quality was assessed using an RNA 6000 Nano kit in a 2100 Bioanalyzer (Agilent Technologies Denmark ApS). RNA was submitted to Novogene Co., Ltd for library preparation and NovaSeq PE150 sequencing. The BBTools package (BBMap - Bushnell B.- sourceforge.net/projects/bbmap) was used for read quality control: bbduk was used for quality trimming with the recommended settings, rRNA removal was done using bbsplit, leaving between 9.26 to 12.1 million reads for each library and optical duplicates were removed using the clumpify function. Post-QC reads were aligned to the reference genomes of *S.islandicus* LAL14/1 and SIRV2gp21 using bowtie2 (Langmead and Salzberg, 2012) and reads for annotated regions in each library were counted with FeatureCounts v2.0.0 (Liao et al., 2014). Differential gene expression analysis was done using the R packages DESeq2 (Love et al., 2014) with a Walden test and EdgeR (Robinson et al., 2010) and genes with a p<= 0.05 were considered as differentially regulated. Functional annotation of the open-reading frames in the *S. islandicus* LAL14/1 genome was achieved by retrieving the COG annotation of the genome from IMG/M (Chen et al., 2017), assigning KO terms with the KOfamKOALA web-server (Aramaki et al., 2020) and matching open-reading frames to the Conserved Domain Database (CDD) (downloaded on October 2019) (Marchler-Bauer et al., 2017) using DELTA-BLAST (Boratyn et al., 2012) (with the options - evalue 0.01, -max_hsps 1 and -max_target_seqs 1). To evaluate gene conservation within the Sulfolobales, OrthoFinder v2.5.4 (Emms and Kelly, 2019) was used to determine orthologous genes and obtain a count-matrix of occurrences in 15 representative Sulfolobales genomes. The results of the above analyses are deposited in the Supplementary Table S2.

### Electrophoresis Mobility Shift Assay (EMSA)

DNA probes were obtained either as complementary oligonucleotides annealed into dsDNA or by PCR amplification of 200 bp fragments corresponding to the upstream region of the indicated genes of *S. islandicus* LAL14/1 (Supplementary Table S8). DNA probes were radiolabeled with [γ-^32^P]-ATP (PerkinElmer) using the T4-polynucleotide kinase (New England Biolabs). For EMSA assays, 30 nM radiolabeled probe and 100 nM protein were incubated in 20 μL binding buffer (10 mM Tris-HCl, pH 8.0, 10 mM HEPES, 1 mM EDTA, 1 mM dithiothreitol, 50 mM KCl, 50 μg/ml bovine serum albumin) with increasing concentrations of cold competitor DNA (ratios 1:1, 1:10, 1:20 and 1:50) for 30 min at 48 ℃ (Guillière et al., 2009). Reactions were then mixed with the 6x loading buffer (0.02% bromophenol blue, 40% glycerol, v/v) and samples were loaded into a non-denaturing 12% acrylamide gel in 0.5X TBE buffer (65 mM Tris pH 7.6, 22.5 mM boric acid and 1.25 mM EDTA). Results were recorded by phosphorimaging with a Typhoon FLA-7000 scanner.

### Prediction of binding-motifs

Enriched motifs present in the 80 bp upstream region of the SiL_0190 or SIRV2 gp21-bound promoters, or the corresponding sequence of the homolog genes of other members of the Sulfolobales (*Sulfolobus acidocaldarius* DSM 639*, Sulfurisphaera tokodaii, Stygiolobus azoricus* DSM 6296*, Metallosphaera hakonensis* DSM 7519 *and Acidianus manzaensis*), were predicted by word-based analysis (k = 6, 7 and 8) using the RSAT pipeline (van Helden et al., 1998; Nguyen et al., 2018).

### Fluorescence microscopy and flow cytometry

Protein expression in triplicate cultures of the *Sulfolobus islandicus* strains harboring constructs for the expression of different RHH-domain factors or pGE2 as a control was induced at OD600 0.05 by the addition of D-arabinose to a concentration of 0.2%. The growth of the cultures was subsequently monitored by measuring absorbance at 600 nm and a 5 ml aliquot was collected at the indicated time points for microscopy and flow cytometry analyses as described previously (Bernander and Poplawski, 1997; Martínez-Alvarez et al., 2017). Briefly, for fluorescence microscopy 1.5 ml samples at OD_600_=0.2 or the equivalent number of cells at each collection point were centrifuged and fixed with 4% paraformaldehyde, followed by a series of washes with PBS (137 mM NaCl, 2.7 mM KCl, 10 mM Na_2_HPO_4_, 2 mM KH_2_PO_4_). Next, each sample was spotted on a poly-L-lysine slide (Sigma-Aldrich), dried and stained with 10 μM 4’,6-diamino-2- phenylindole (DAPI) (Invitrogen). After another round of PBS wash, the slides were covered with mounting medium (78% glycerol, 1X PBS, 0.2% polyvinylpyrrolidone) and stored at 4°C until analysis. The slides were analyzed with the Nikon Eclipse Ti inverted microscope at 100X magnification and images were collected using the software NIS-Element (RRID: SCR_014329). For flow cytometry analyses, a volume of 300 μl of culture at OD_600_=0.08 or the equivalent number of cells were fixated in ethanol (final concentration of 70%) and incubated overnight. The fixed cells were centrifuged at 2,800 rpm for 20 min and resuspended in 1 mL wash buffer (10 mM Tris pH 7.5, 10 mM MgCl2) and collected again by centrifugation. Finally, the cell pellets were resuspend in 140 μL of staining solution (the washing buffer plus 40 μg/mL ethidium bromide (Thermo-Scientific) and 100 μg/mL mithramycin A (Apollo Scientific), and stained for at least 20 min on ice. Stained cells were analyzed in an Apogee A10 Bryte (Apogee Flow Systems) and a total of 35 000 events were collected for each sample. Two parameters were measured: FSC (forward scattered light) and SSC (side scattered light). The results were plotted and analyzed with the FlowJo™ v10.8.1 and Flowing Software 2, the ratio of normal cells vs. enlarged cells was calculated for each factor.

### RNA extraction and cDNA synthesis

Samples (corresponding to approximately 1 × 10^9^ cells) were collected by centrifugation and stored in 1 mL TRI reagent (Sigma Aldrich) at −80°C. Total RNA was extracted using the TRI reagent following the instructions by the manufacturer. RNA pellets were washed and dissolved in nuclease-free water; RNA integrity verified by gel electrophoresis and concentration was measured using a Nanodrop 1000 spectrophotometer.

RNA samples were first treated at 37°C for 30 min with RNase-free DNase I (Invitrogen) in 50 μl reactions containing about 5μg RNA and 1 unit DNase I. The treatment was stopped by adding 2 μl EDTA to a final concentration of 5 mM, followed by incubation at 75°C for 10 min. A 3 μl-treated RNA aliquote was reverse transcribed in 10 μl reaction using SuperScript II reverse transcriptase (Invitrogen) according to manufactory’s instruction. As a control for possible contamination from genomic or viral DNA, another 3 μl RNA was processed in the same way except that no reverse transcriptase was added.

### Quantitative real-time PCR

The reactions were performed in 10 μL mixtures containing 5 μL 2x SYBR Green Supermix (Bio-Rad), 0.5 mM primers and about 1 ng cDNA. Separate reactions were prepared for detection of reference genes (Supplementary Table S7). The mixtures were prepared in 96-well microtiter PCR plates (Bio-Rad Laboratories), sealed with an adhesive cover (Bio-Rad Laboratories) and amplified on the CFX96 Real-Time Detection System (Bio-Rad Laboratories) with the following program: denaturing at 95℃ for 30s, 40 cycles of 95℃ for 5s and 56℃ for 30s. Additionally, a non-template control (NTC) was used to check the possibility of unspecific amplification products and contamination. 16S rRNA was used as the reference and *tbp*, a housekeeping gene encoding the TATA-binding protein, was used as a control.

### Sequences, identification of homologous proteins and phylogenetic analysis

A manually curated list of genomes from 200 archaeal viruses deposited in NCBI (downloaded on February 2020) and including proviral genomes of thaumarchaeal viruses (Krupovic et al., 2019) was used to construct a database containing 11 120 proteins hereby referred to as archaeaVir. A list of genomic accession numbers and identifiers is provided in Table S1. Functional annotation of the proteins in the archaeaVir database was assigned by using the program hmmsearch of the HMMER suite (Eddy, 2011) to match the sequences to the VOG (Grazziotin et al., 2017) and pfam (El-Gebali et al., 2019) databases, and by using DELTA-BLAST (Boratyn et al., 2012) to match sequences to the Conserved Domains Database (CDD) (Marchler-Bauer et al., 2017). Proteins annotated to have an RHH-domain were retrieved and used as queries for the identification of homologous proteins in the archaeaVir database by using PSI-BLAST (Altschul et al., 1997)(Altschul et al 1997) with a relaxed e-value of 0.1 to identify divergent homologs. If two or more queries had overlapping matches, they were merged into the same cluster of homologous proteins. Cellular homologs of gp21 were initially identified through a protein BLAST search to the genome of *S. islandicus* LAL14/1 (evalue 0.1), which retrieved 3 hits: *SIL_RS00945, SiL_RS14755* and *SIL_RS07870*. A PSI-BLAST iteration of the search added one more hit corresponding to the conserved gene *SIL_RS00595*. A set of proteins from 15 members of the Sulfolobales order was then used to perform a protein BLAST search (evalue 0.001) to identify the corresponding homolog in every genome, keeping the match with the highest score or the match that has the same genomic context as in the query. Synteny at the genomic level was evaluated using SyntTax (Oberto, 2013). Sequences of the proteins containing more than two RHH domains, commonly found in homologs in the *Bicaudaviridae* viral family, were split into their individual domains for a better comparison. To construct a phylogenetic tree of the SIRV2 gp21 orthologs, viral and cellular sequences were aligned using MUSCLE (Edgar, 2004), followed by maximum likelihood phylogenetic analysis with FastTree 2 (v2.1.3) (Price et al., 2010) using an LG+CAT model and a Gamma20-based likelihood (option-gamma). The resulting tree was visualized with FigTree v1.4.4 (http://tree.bio.ed.ac.uk/software/figtree/).

### Model of SIRV2 gp21 and SiL_0190

The structures of SiL_0190, SIRV2 gp21 and other members of the family were predicted using AlphaFold2 (Jumper et al., 2021). Alignment of the predicted structures to a DNA molecule was done by superimposing the models to the crystal structure of the ArtG transcriptional regulator of *Staphylococcus aureus* bound to its target DNA (PDB: 3GXQ).

## Acknowledgements

We acknowledge de Biocomputing Core Facility of the Department of Biology, University of Copenhagen for access to resources used for bioinformatic analyses. This work was supported by Independent Research Fund Denmark/Natural Sciences (DFF-0135-00402) and Novo Nordisk Foundation/Hallas-Moeller Ascending Investigator Grant (NNF17OC0031154) to X.P..

## Author contributions

X.L, L.M.-A and X.P conceived the experiments. X.L, C.L.-M and L.M.-A performed the experiments. X.L, L.M.-A and X.P wrote the paper. All authors revised and approved the manuscript.

---

A Prophage-Encoded Small RNA Controls Metabolism and Cell Division in Escherichia coli - PubMed.

**Figure S1.**
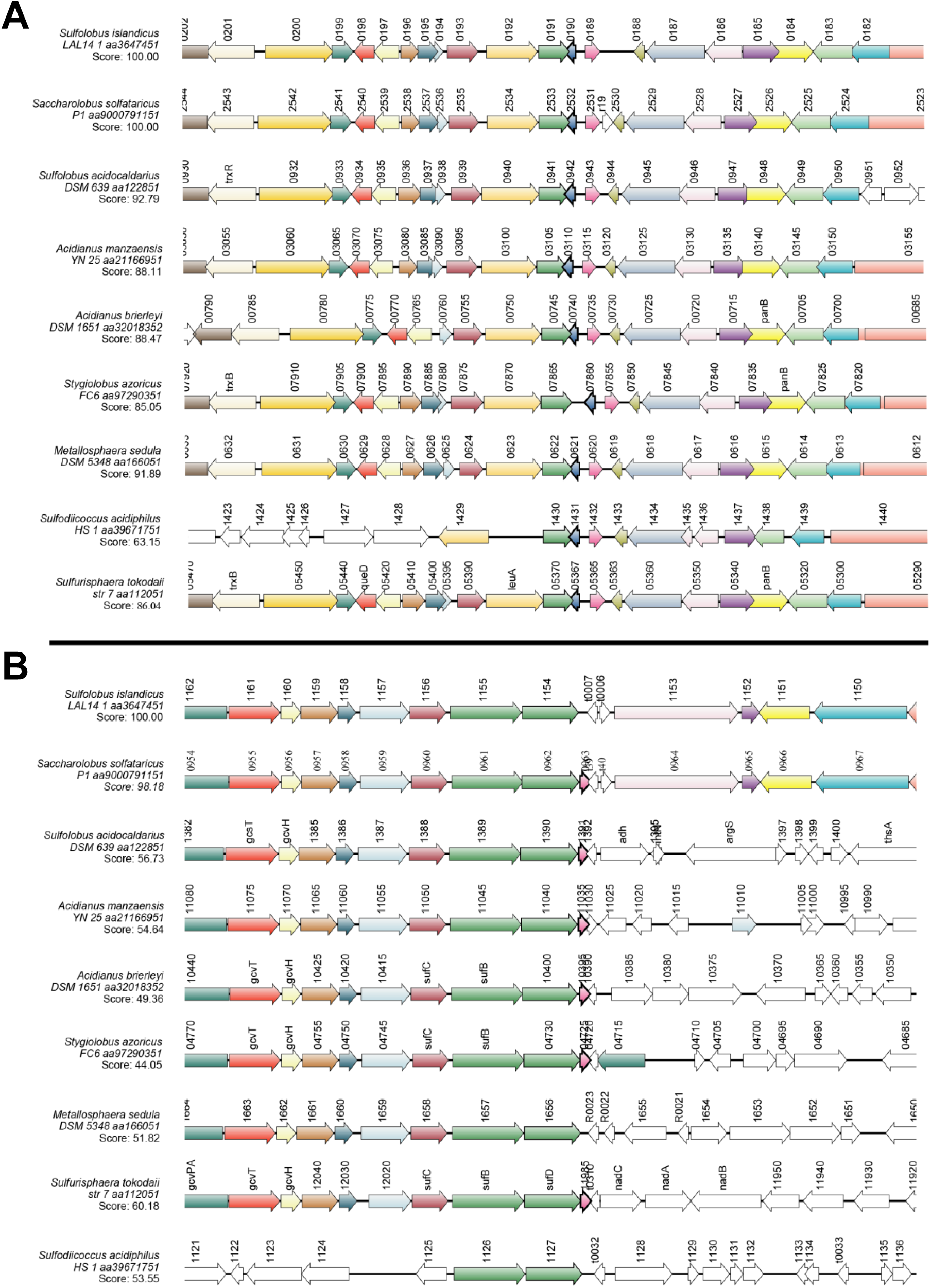
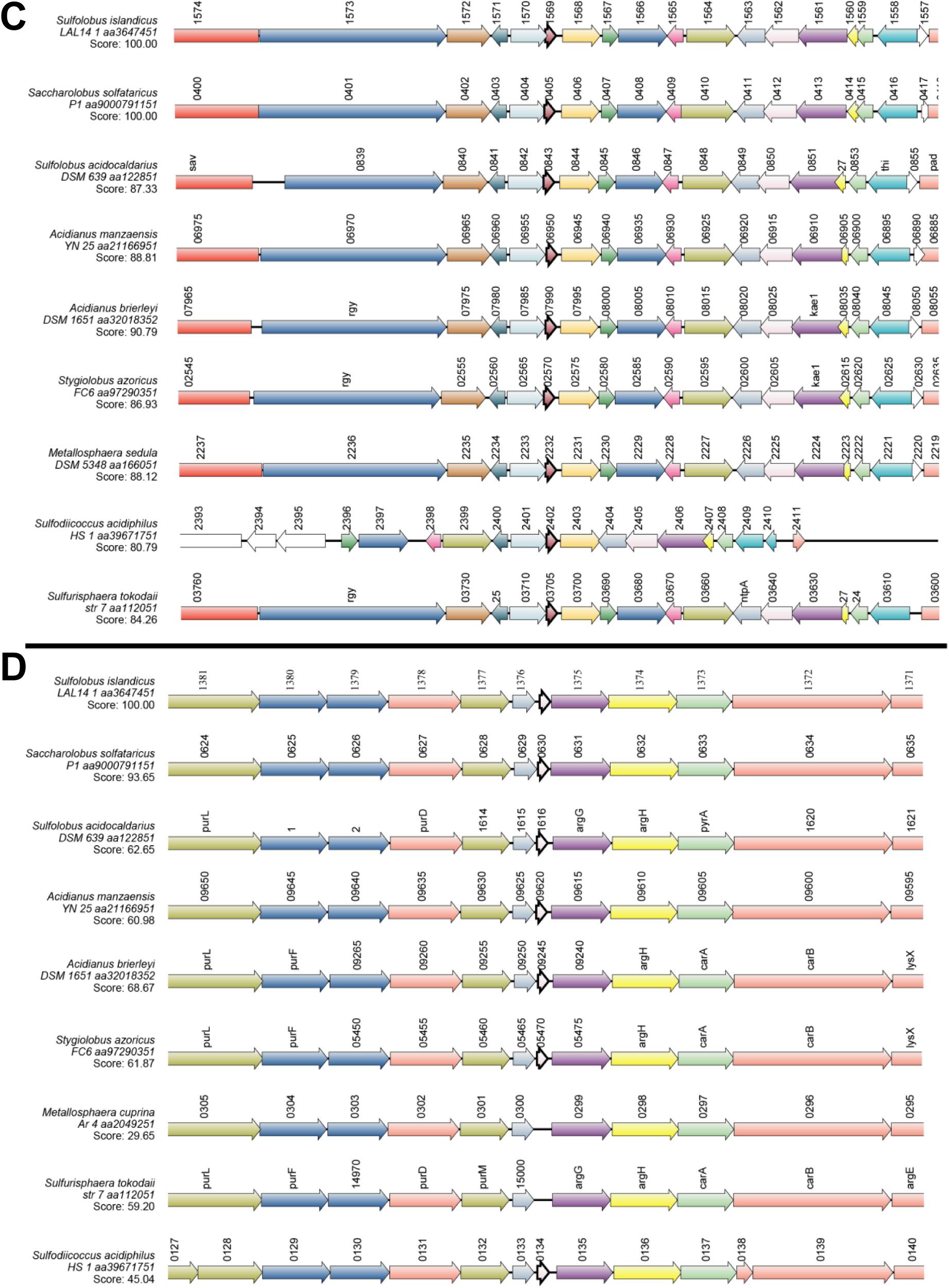
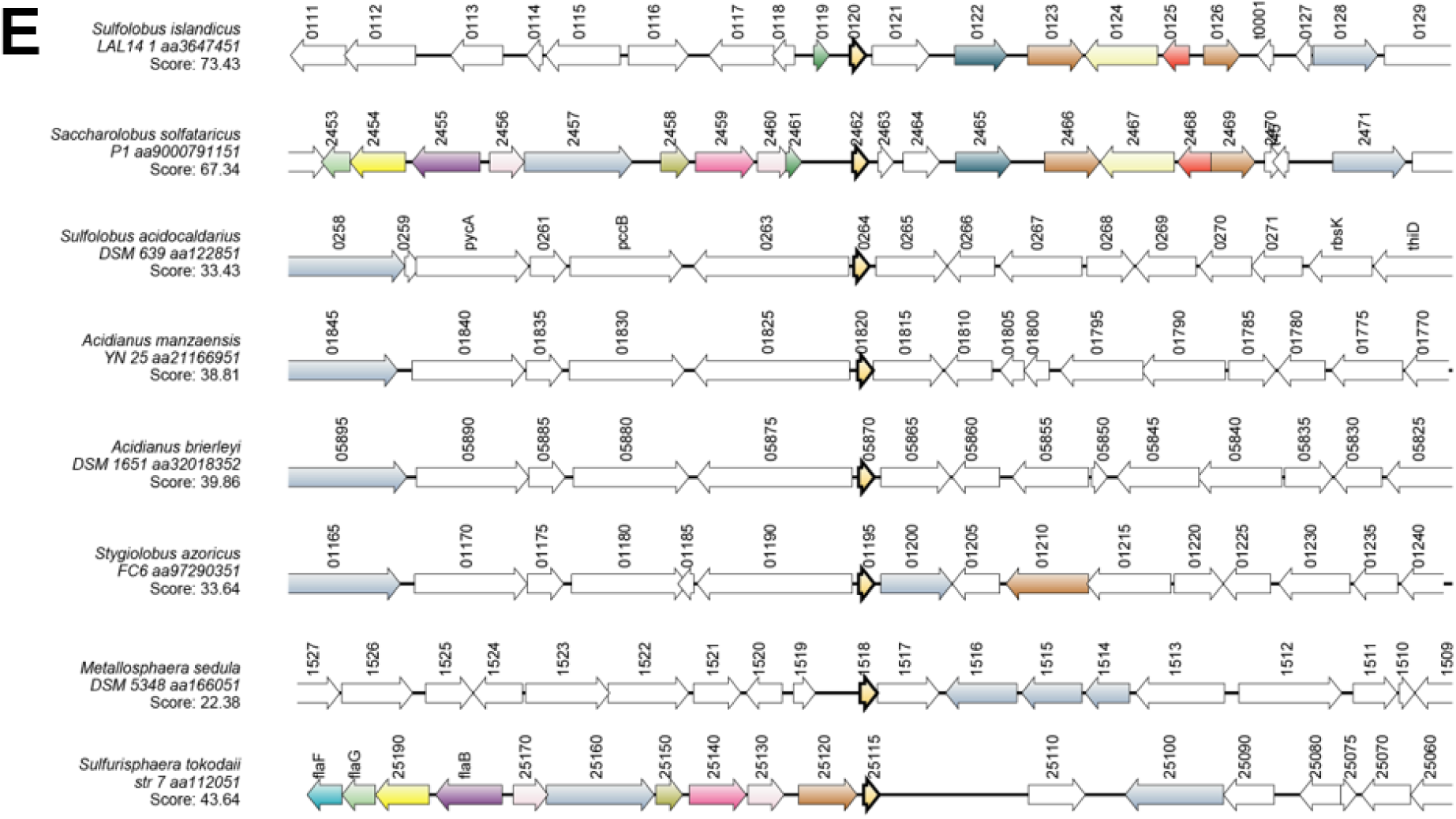
Genomic neighborhood of the cellular homologs of SIRV2gp21 in different members of the Sulfolobales. A) SiL_0190, B) SiL_RS14755, C) SiL_1569, D) SiRe_1373 homolog in LAL14/1 (1 286 819 to 1 286 682 bp), E) SiL_0120. Results obtained using SyntTax (Oberto et al. 2013). The query proteins are outlined in bold and the legend indicates the BLAST score of paralogs to the LAL14/1 query. Paralogs are indicated with the same color across genomes.

**Figure S2.**
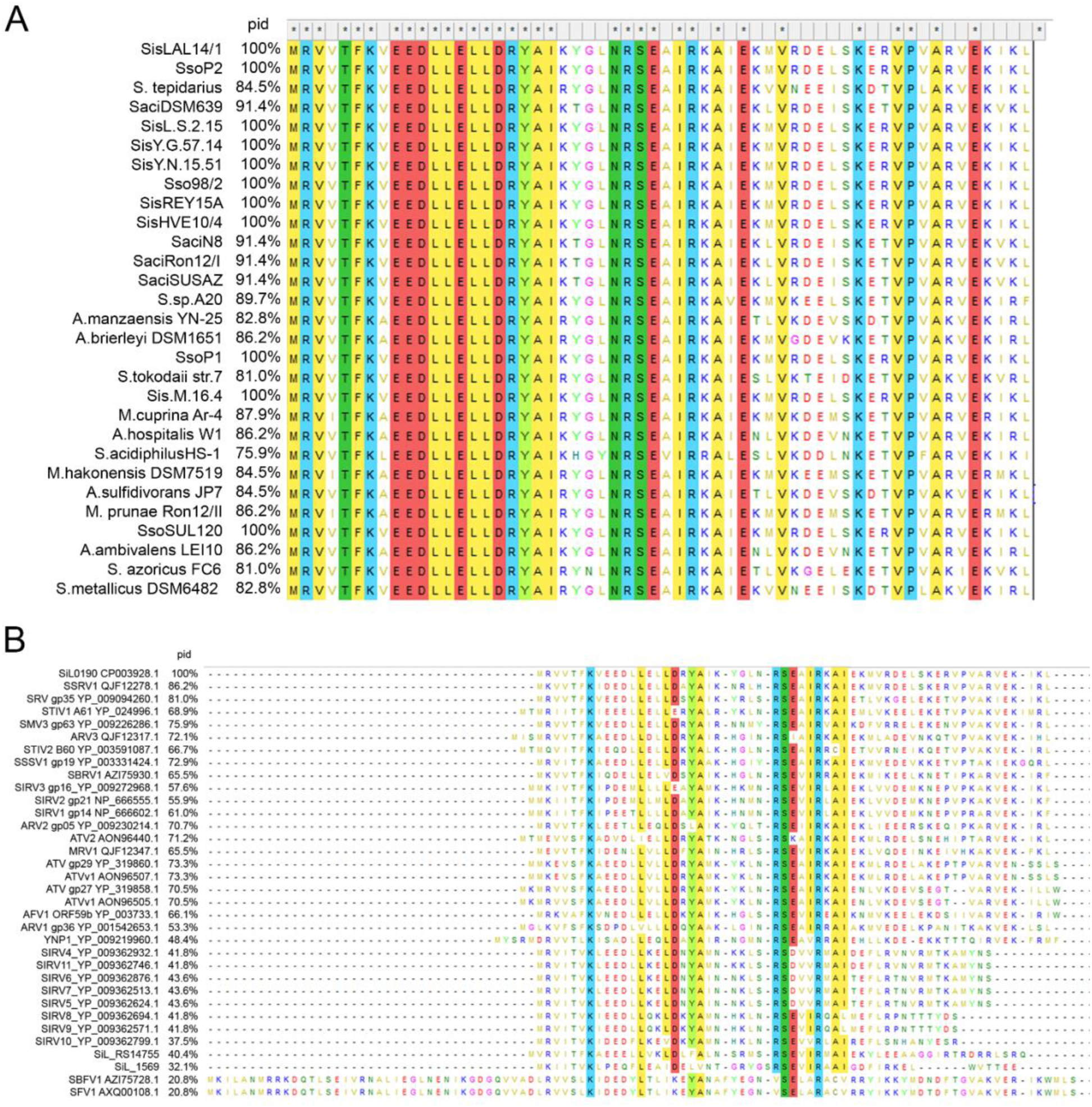
Sequence alignment of members of the SiL0190 clade of RHH proteins. **A)** Cellular homologs in 29 members of the Sulfolobales. **B)** Viral homologs aligned to SiL_0190. The percentage identity to SiL_0190 is indicated on the left. Sequences were aligned using MEGAX with the MUSCLE algorithm (Xie et al. 2008) (Edgar et al. 2004).

**Figure S3.**
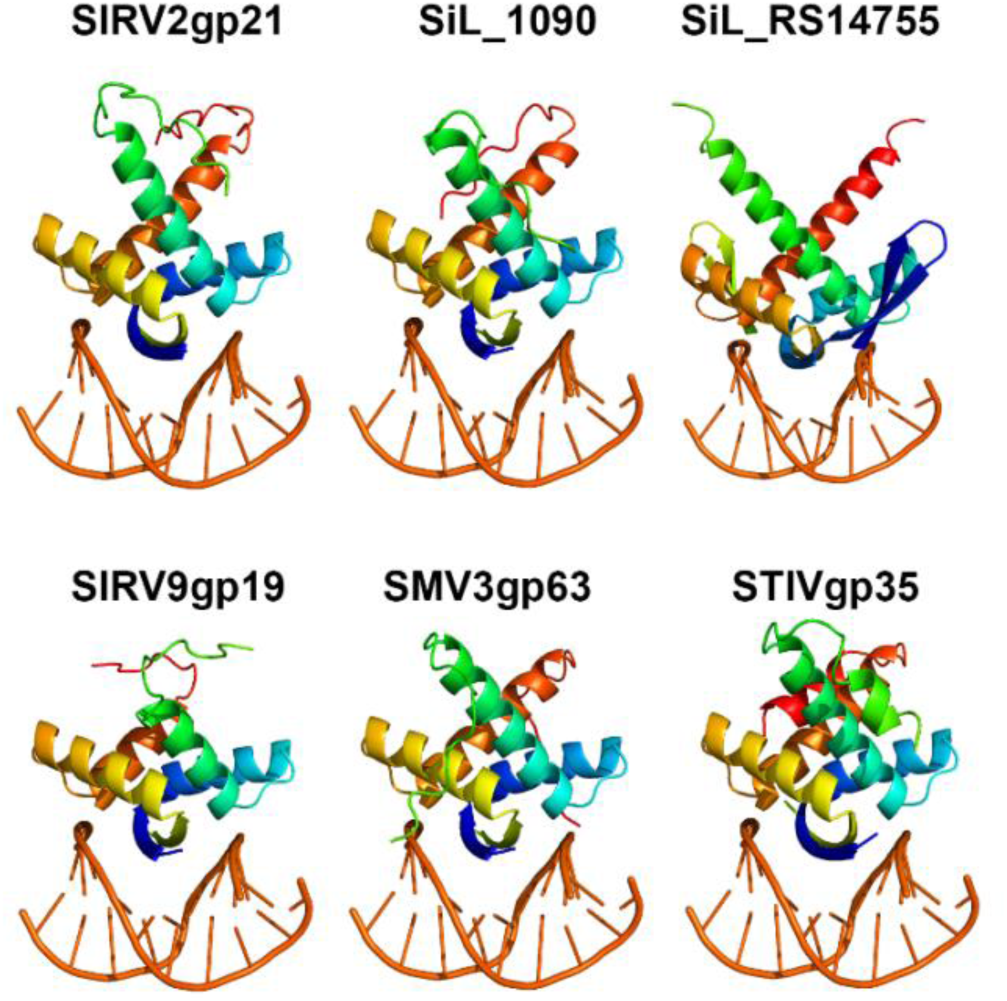
Structural models of members of the SIRV2 gp21 protein family. Proteins were modeled with AlphaFold2 (Jumper et al. 2021) and superimposed to the structure of the RHH-domain ArtA regulator with its target DNA (PDB: 3GXQ). All structures are predicted to dimerize.

**Figure S4.**
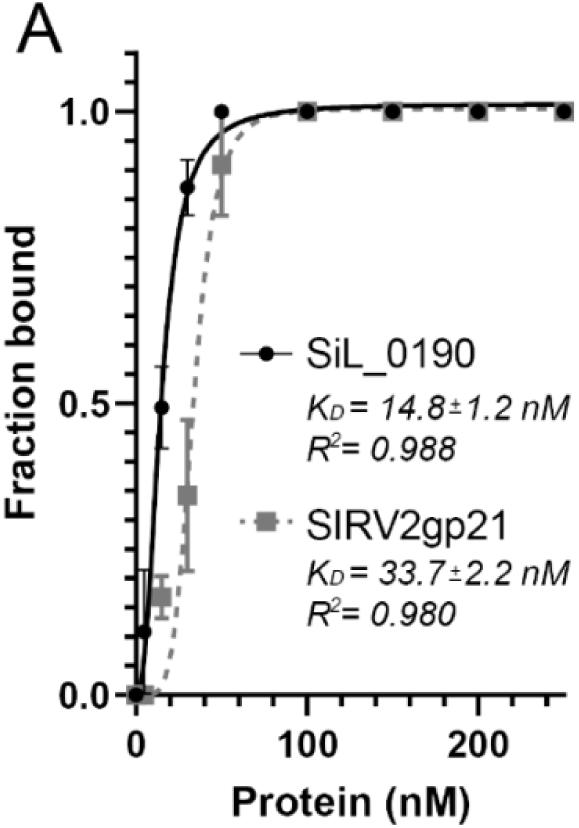
Apparent DNA binding affinity of SiL_0190 and SIRV2 gp21 in the presence of competitor DNA. **A)** The binding affinity (K_D_) of each protein was determined with EMSA assays by mixing 15 ng labeled *cdvA* promoter and 1500 ng of competitor DNA with increasing concentrations of the corresponding protein. Graph show the average and standard deviation of three independent replicates.

**Figure S5.**
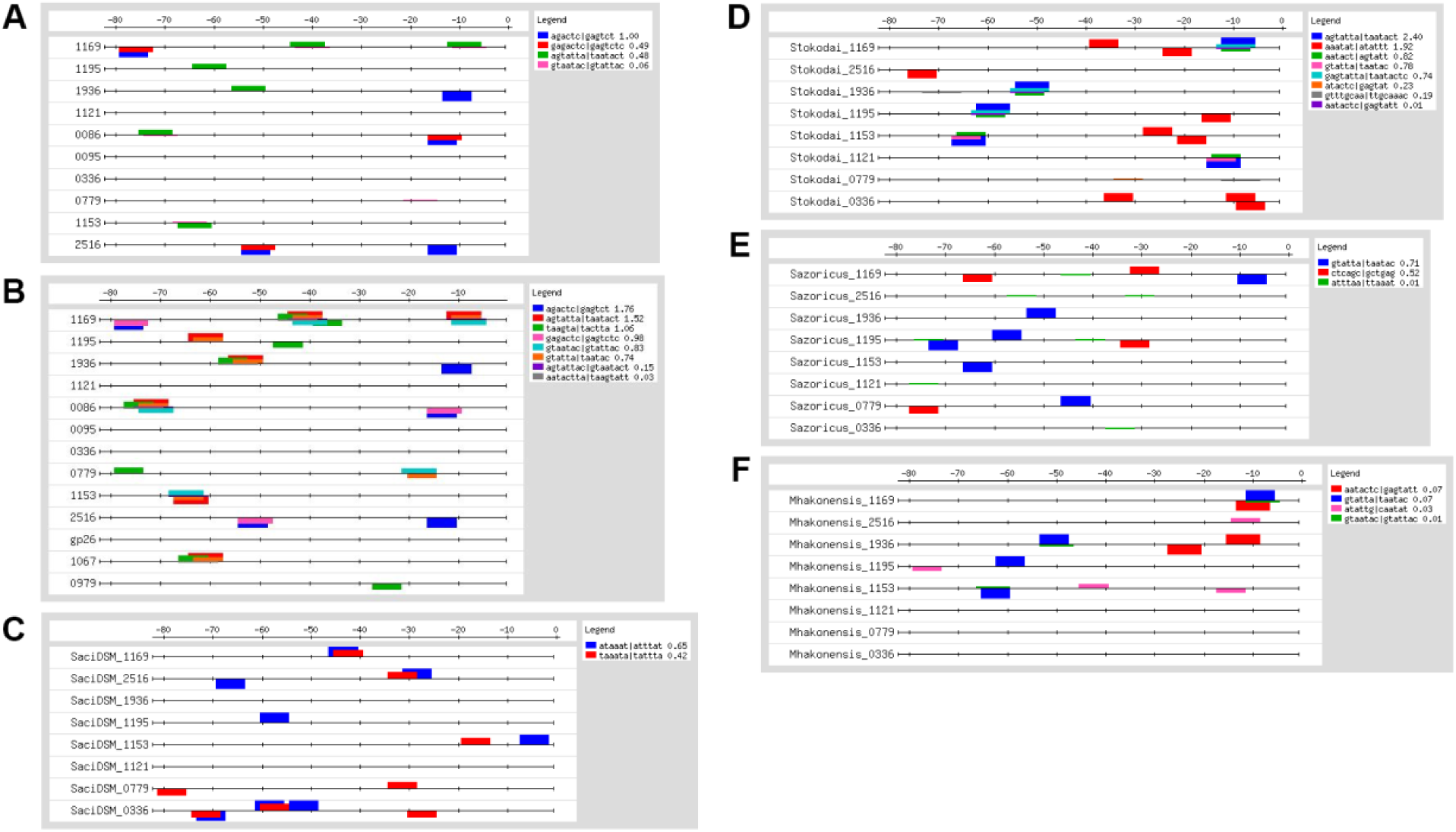
Prediction of enriched oligonucleotides in the promoters of the SiL_0190 and SIRV2 gp21 target genes. Analysis of oligonucleotide frequencies was done using the RSAT oligo-analysis tool with the regulons (80 nucleotides upstream of the gene start codon) bound by SiL_0190 **(A)** and SIRV2 gp21 **(B)** in the EMSA assays. **(C—F)** The search for enriched oligonucleotide sequences was repeated in different members of the Sulfolobales by using the promoter region of the homolog genes of the SiL_0190 targets in the corresponding strain. **C)** *Sulfolobus acidocaldarius* DSM 639, **D)** *Sulfurisphaera tokodaii*, **E)** *Stygiolobus azoricus* and **F)** *Metallosphaera hakonensis*.

**Figure S6.**
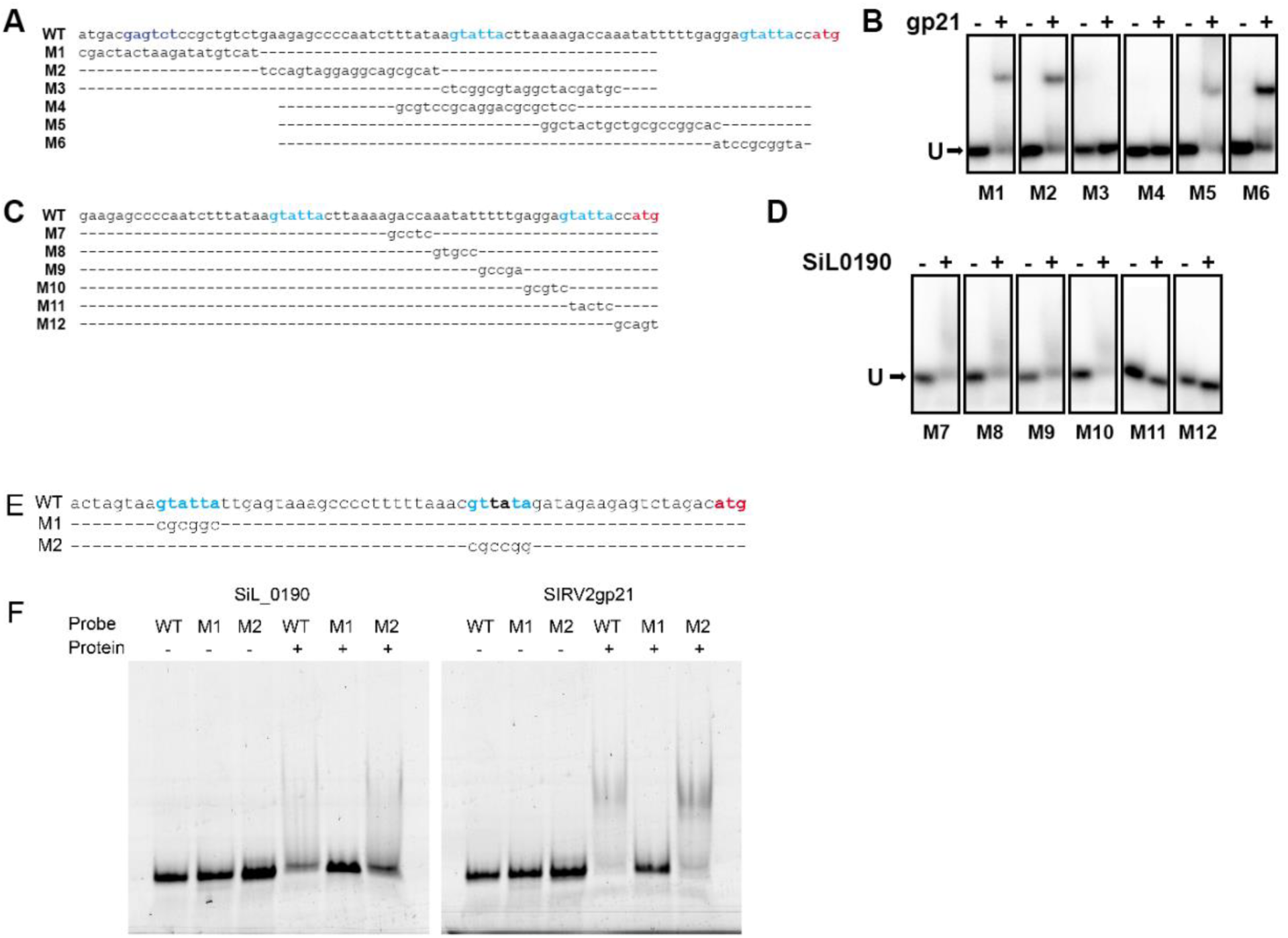
SiL_0190 and SIRV2 gp21 bind to the AGTATTA motif. **A)** Upstream region of the *cdvA* gene of *S.islandicus* LAL14/1 and variants with mutations across the sequence (M1-M3). **B)** EMSA assay of SIRV2 gp21 and the promoter variants in panel A. While mutation of the GAGTCT motif (located between −76 and −70) does not impair binding, mutation of the AGTATTA motif (−35 to −41) abrogates binding of SIRV2 gp21. **C)** Upstream region of the *cdvA* gene and variants M7-M12 with 5 base pairs mutations. **D)** EMSA assay of SiL_0190 to the promoter variants in panel D. **E)** Promoter of the *sil_1936* gene and variants with mutations in the DNA-binding motif (M1) or in a degenerate motif sequence (M2). **F)** EMSA assay of SiL_0190 (left) or SIRV2 gp21 (right) with the promoter variants in panel E. The AGTATTA motifs are indicated in light blue and the GAGTCT motif is highlighted in dark blue. The start codon of the genes is highlighted in red. Oligonucleotides used for EMSA assays in this figure are described in Supplementary Table S8.

**Figure S7.**
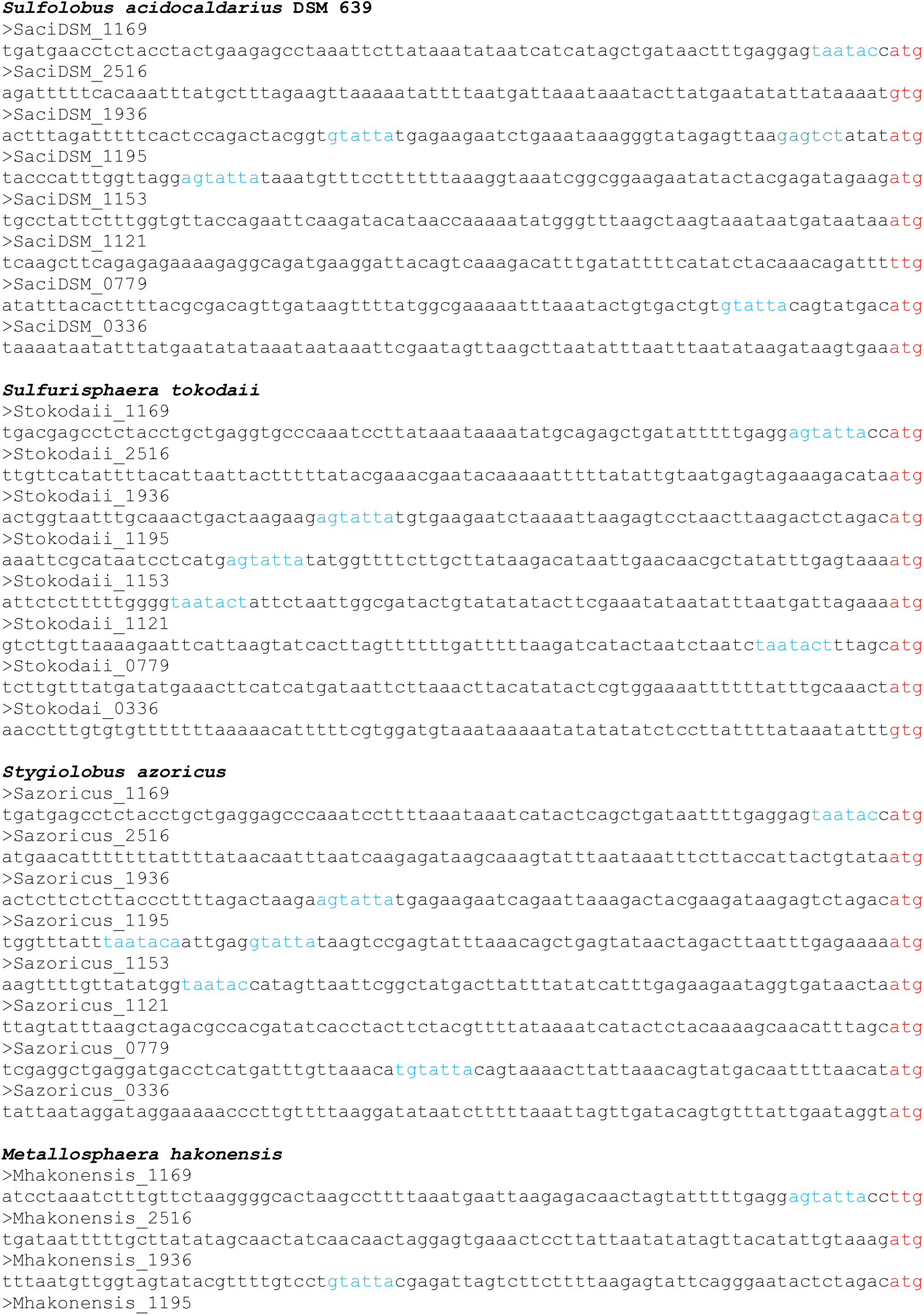
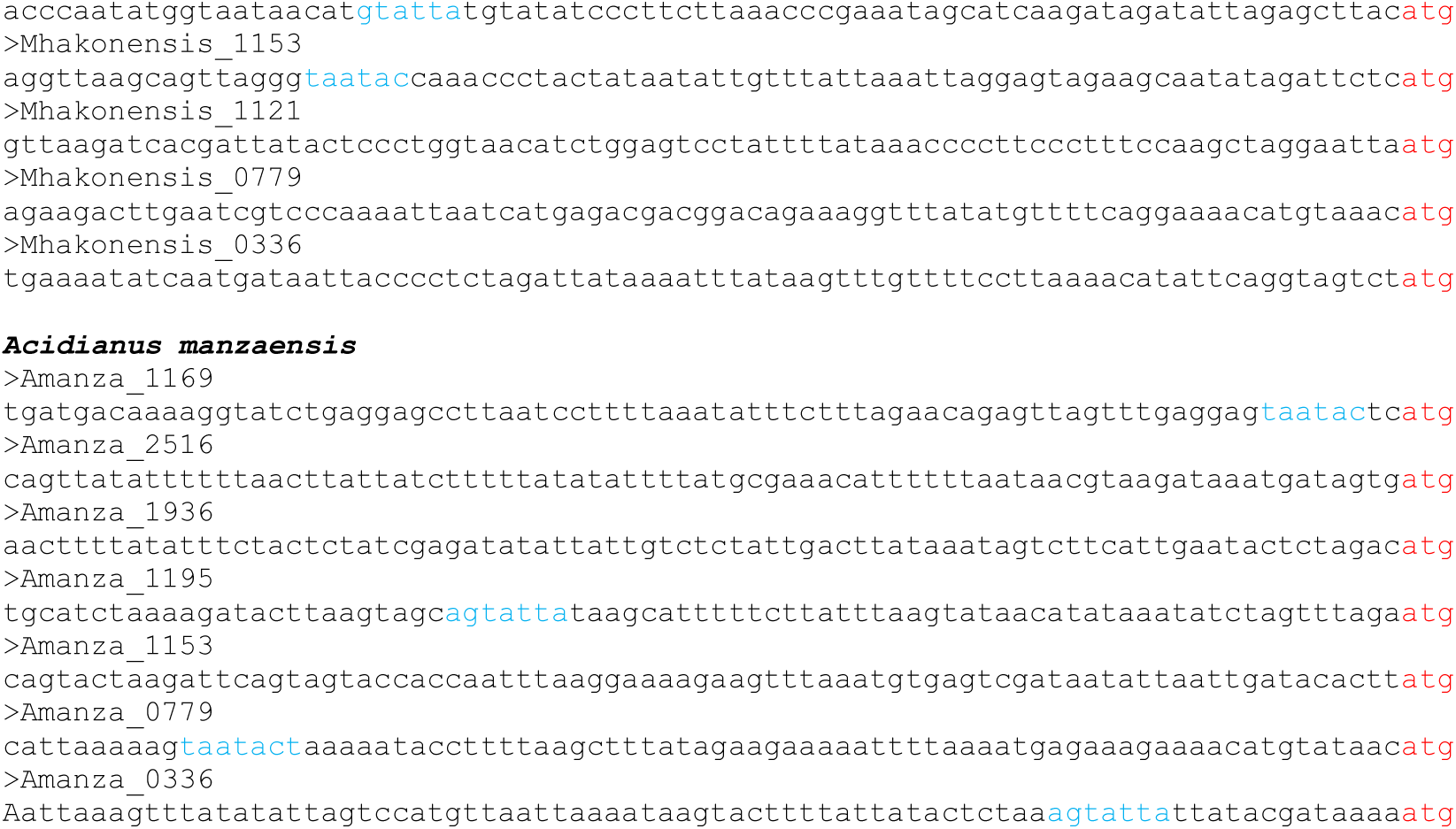
The motif AGTATTA|TAATACT is present in the promoters of the homologs of the SiL_0190 target genes in different Sulfolobales. Upstream sequences of the homologs of *sil_1169, sil_1195, sil_1935-1936, sil_1153, sil_0336, sil_0779* and *sil_2516* are shown for four different members of the Sulfolobales order. The start codon of the gene is highlighted in red and occurrences of the AGTATTA|TAATACT motif (or the seed GTATTA|TAATAC) are highlighted in blue.

**Fig S8.**
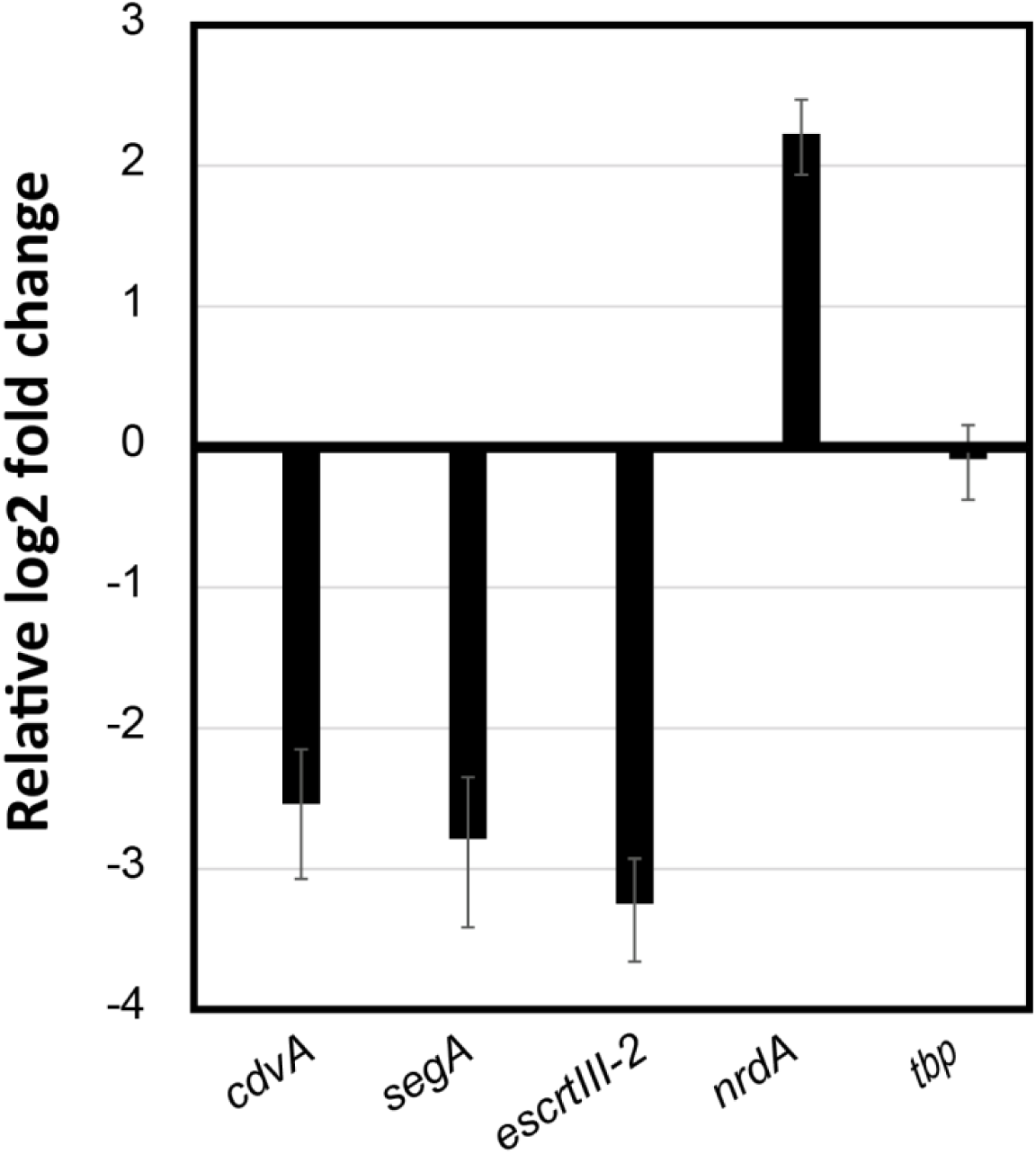
Expression of SiL_0190 target-genes after the induction of SiL_0190. *S. islandicus*. Δarrays cultures harboring a plasmid for the inducible expression of SiL_0190 were sampled and analyzed by RT-qPCR at 6 hours post-induction with arabinose. 16S rRNA was used as the normalization reference and *tbp* (TATA-binding protein) as the control. The graph shows the average and standard deviation of three biological replicates

**Figure S9.**
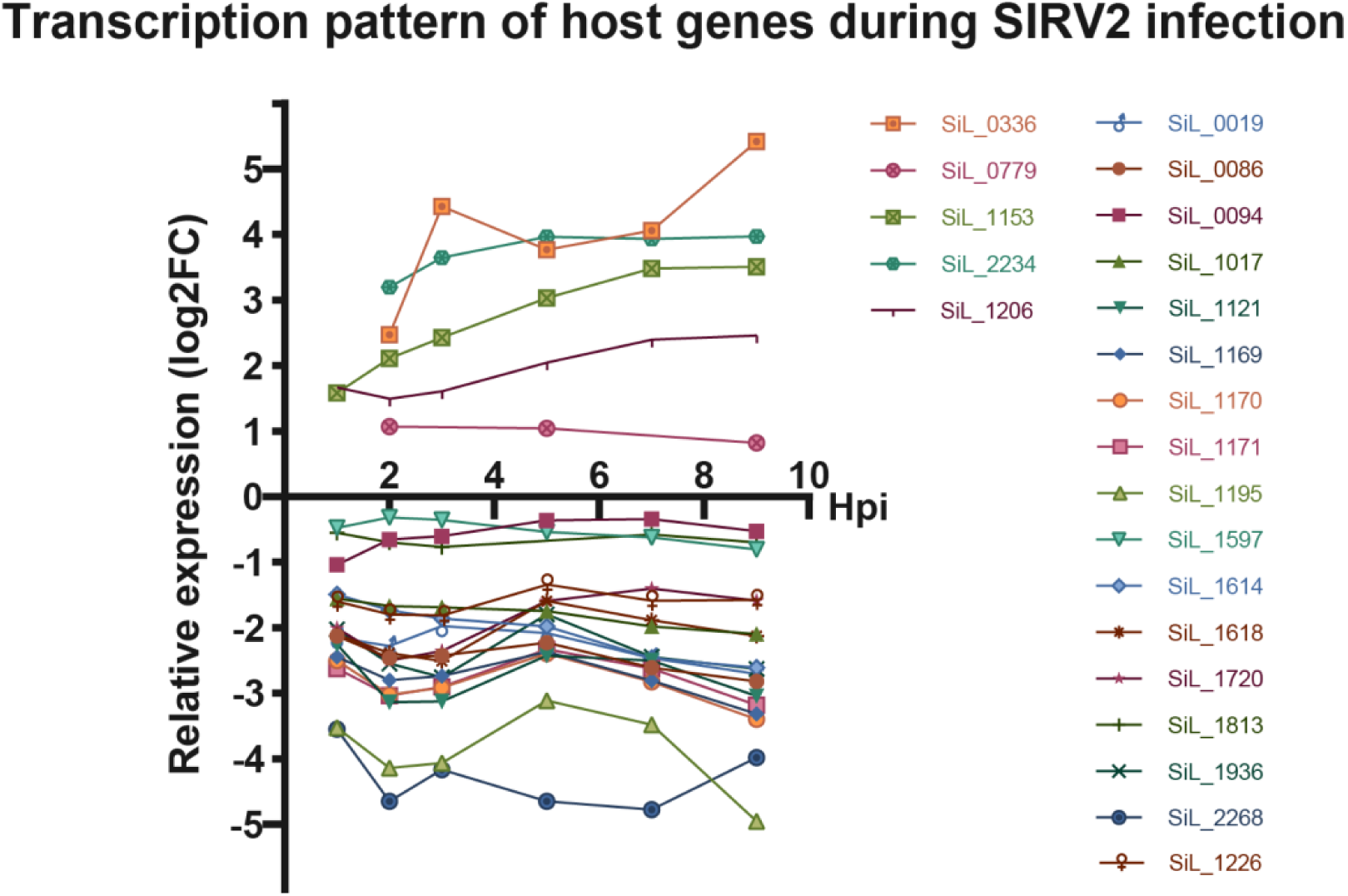
Transcriptional pattern of selected cellular genes in SIRV2-infected cells. The data represents the relative log2 fold change (log2FC) in the transcription level of the indicated genes in infected *S. islandicus* LAL14/1 cells with respect to uninfected cultures. Hpi – hours post-infection. Graphical representation of the transcriptomic data by Quax et al. (2013). Oligonucleotides used for RT-qPCR analyses are listed in Supplementary Table S9.

